# Information-Distilled Generative Label-Free Morphological Profiling Encodes Cellular Heterogeneity

**DOI:** 10.1101/2023.11.06.565732

**Authors:** Michelle C.K. Lo, Dickson M. D. Siu, Kelvin C. M. Lee, Justin S. J. Wong, Maximus C.F. Yeung, Michael K.Y. Hsin, James C.M. Ho, Kevin K. Tsia

## Abstract

Image-based cytometry faces constant challenges due to technical variations arising from different experimental batches and conditions, such as differences in instrument configurations or image acquisition protocols, impeding genuine biological interpretation of cell morphology. Existing solutions, often necessitating extensive pre-existing data knowledge or control samples across batches, have proved limited, especially with complex cell image data. To overcome this, we introduce *Cyto-Morphology Adversarial Distillation* (CytoMAD), a self-supervised multi-task learning strategy that distills biologically relevant cellular morphological information from batch variations, enabling integrated analysis across multiple data batches without complex data assumptions or extensive manual annotation. Unique to CytoMAD is its “morphology distillation”, symbiotically paired with deep-learning image-contrast translation - offering additional interpretable insights into the label-free morphological profiles. We demonstrate the versatile efficacy of CytoMAD in augmenting the power of biophysical imaging cytometry. It allows integrated label-free classification of different human lung cancer cell types and accurately recapitulates their progressive drug responses, even when trained without the drug concentration information. We also applied CytoMAD to jointly analyze tumor biopsies across different non-small-cell lung cancer patients’ and reveal previously unexplored biophysical cellular heterogeneity, linked to epithelial-mesenchymal plasticity, that standard fluorescence markers overlook. CytoMAD holds promises to substantiate the wide adoption of biophysical cytometry for cost-effective diagnostic and screening applications.

## 1. Introduction

Recent breakthroughs in label-free single-cell imaging have highlighted the significance of biophysical cytometry in understanding functional cellular heterogeneity in complex biological systems [1]. Supercharged by their increasing throughput and content, systematic profiling of biophysical cell morphology, which was once inconceivable, is now feasible to probe subtle differences in cell mass, shape, size, and biophysical/mechanical properties across different cell types/states or responses to various chemical and genetic perturbations [2–8]. These imaging technologies are gaining traction in pharmaceutical industries and life science laboratories, providing critical mechanistic insights into cellular functions that may be concealed in molecular assays [9, 10].

However, cellular imaging experiments are notoriously complicated by batch effects that result from differences in imaging system configurations, image acquisition protocols, and experimental conditions. Specifically, in microfluidic imaging flow cytometry, batch-to-batch variations manifest from various sources, encompassing (1) the optical system factors such as power instability of the laser source and the noise originated from photodetection and signal amplification in the system; (2) the microfluidic aspects associated with the variations in the microfluidic chip fabrication quality that could potentially result in the image distortion/aberration. These unwanted technical variations are no exception in biophysical imaging cytometry, where they obscure genuine biological signals and hinder robust integrated data analysis across multiple datasets. Consequently, minimizing batch effect is of paramount importance, yet challenging, for improving data reproducibility and accurately capturing biological information [11].

Batch normalization is a widely used strategy for image-based batch correction [12, 13]. However, current methods, including those based on machine learning techniques, face several limitations that can hinder their effectiveness in correcting batch effects. Firstly, many methods require a priori knowledge or assumptions about the statistical distributions within each batch to align the data across different batches. Secondly, some methods necessitate the presence of a common control sample (e.g., a negative control) across all batches for normalization. In parallel, a few methods have been developed to address batch effect correction in single-cell omics analysis [14, 15]. However, the inherent differences between 1D sequencing data and 2D (or even 3D) images, together with the significant diversity and complexity of biological image data structures, limit the direct applicability of these methods in image-based cellular/tissue analysis. Furthermore, these approaches typically fall short in generating batch-effect-free images directly for downstream analysis. Consequently, disentangling batch effects from biological image datasets remains a challenging task in imaging cytometry and is more complex compared to traditional omics data. This underscores the need to develop more effective batch effect correction methods tailored to image-based data.

Here, we introduce a new generative deep-learning pipeline, *cyto-morphology adversarial distillation* (CytoMAD). This approach not only enables batch-effect correction, but also allows simultaneous label-free image contrast translation to reveal additional cellular information. In this work, we primarily focus on label-free imaging modalities (i.e., bright-field (BF) to quantitative phase image (QPI) translation) due to their growing significance in biology, as they uncover biophysical and mechanical properties of cells that underpin cell functions, which are not always discernible from fluorescence counterparts [16–18].

In contrast to previous deep-learning batch correction approaches [13, 19, 20] or image translation approach [21–25], CytoMAD offers three key unique attributes: (1) flexibility in modeling complex, non-linear data distributions, enabling correction of various batch effects without distributional assumptions; (2) accurate generation of QPI that is applicable to batch effect correction by learning to translate (augment) images across batches while conserving the biological content; (3) it simultaneously allows self-supervised batch-corrected morphological profiles for integrated downstream analysis.

We demonstrate CytoMAD’s diverse capabilities in various applications, including accurate joint analysis across multiple batches for label-free classification of human lung cell types, functional drug-treatment assays for morphological changes in response to a panel of drugs at different concentrations (even the drug concentration is excluded in training), and integrated biophysical cellular analysis of tumor biopsies from multiple non-small cell lung cancer (NSCLC) patients. Our results showcase CytoMAD’s versatility and utility for a wide array of applications in cell biology and biomedical research.

## 2. Results

### 2.1 Overall Framework of CytoMAD

CytoMAD is a multi-task, generative deep-learning model designed to address batch effects in imaging cytometry, with self-supervised learning characteristics. Specifically, its capability to help separate biological information from technical confounders is infused in the process image contrast translation (i.e., QPI outputs predicted from BF images) as well as representation learning (i.e. generation of morphological features/profiles) (Figure 1a). Hence, it allows robust integrated analyses across multiple data batches in the downstream, including joint-dataset classification, biophysical marker discovery and morphological profile interpretation (Figure 1b).

**Figure 1.**
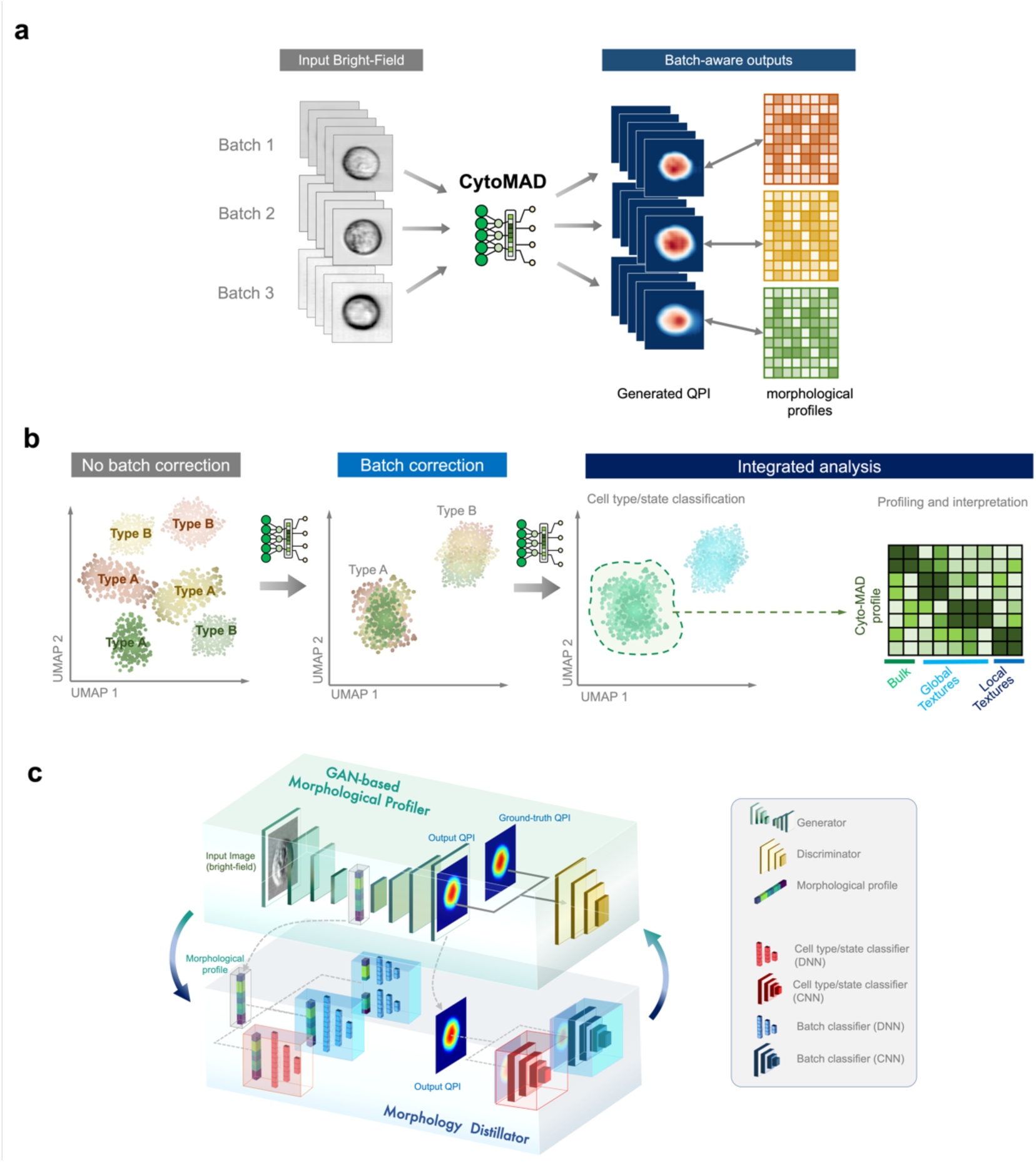
Main framework of CytoMAD. **(a) Inputs and outputs of CytoMAD.** CytoMAD takes in brightfield (BF) cell images from multiple batches as model input. It enables robust image contrast conversion (from BF to QPI) and distills underlying biologically relevant morphological profiles (i.e., CytoMAD profile) from batch distortions. It thus allows robust “batch-aware” downstream morphological profiling and analysis. **(b) General workflow of CytoMAD**. Without CytoMAD, the batch effect confounds the downstream analysis (e.g. classification of cell type A and B from three datasets/batches). CytoMAD corrects the batch effect and thus allows integrated analysis (e.g., classification, profiling and interpretation of the CytoMAD profiles). **(c) Deep-learning architecture of CytoMAD**. The model comprises of two main parts, the GAN-based morphological profiler and the morphology distillator. The GAN-based backbone takes in BF and converts it into QPI output. The discriminator classifies the CytoMAD output from the ground truth and serves as a feedback mechanism to achieve accurate image contrast conversion. The morphology distillator introduces the self-supervised batch-aware characteristic to CytoMAD through a set of classification networks. The classification networks comprise of batch classifiers and cell type/state classifiers, altogether dynamically enabling disentanglement of batch variations from biological information at both the bottleneck morphological features (CytoMAD profile) and output images.

The model consists of two interconnected components: (1) the batch-aware generative-adversarial-network-based (GAN-based) image generation and (2) the morphology distillator (Figure 1c) (See the details in **Methods**). The GAN-based backbone in CytoMAD facilitates image generation and contrast translation for augmented cellular information. Prior to implementing the morphology distillator, the backbone model is pre-trained and guided by a feedback mechanism to achieve accurate image contrast conversion from bright field (BF) to quantitative phase images (QPI). Note that QPI formation bypasses the need for common complex optical QPI instrumentation (e.g., interferometric/holographic modules and multiplexed detection modules) and computationally-intensive QPI reconstruction algorithms [26]. When image contrast translation is not required, a convolutional autoencoder architecture can be used by equalizing the input and output target image of this GAN-based backbone. From this pretrained GAN-based backbone, the resultant morphological features at the GAN’s bottleneck layer (denoted as *no-CytoMAD-profile* at this stage) and the output QPI (denoted as *no-CytoMAD-images* at this stage) would serve as the inputs for training the morphology distillator in the next stage. As the no-CytoMAD-profile is derived from the input image and subsequently utilized in reconstructing the output image, it effectively encodes information from both BF and QPI. Here single cell images are captured by a multi-modal imaging flow cytometer, called multi-ATOM [2, 4, 7, 27], operating at a high imaging throughput of ∼10,000’s cells/sec (See the system details in **Methods**).

CytoMAD distinguishes itself from the common generative model by integrating a self-supervised morphology distillator. This component consists of a set of classification networks, including batch classifiers and cell type/state classifiers (Figure 1c). Both types of classifers work symbiotically to disentangle batch variations from biological information (See the training strategy in **Methods**). The classifier duo is pre-trained based on *no-CytoMAD-profiles* and *no-CytoMAD-images* to identify the batch and cell-type information. The self-supervised nature of the model comes from using image translation and batch classification as pretext tasks, allowing the model to learn meaningful representations without extensive manual annotations. To suppress batch distortion and enhance biological information at the phenotypic features and cellular images with minimal alterations, these classifiers are implemented at both the bottleneck region and the image output of the GAN-based backbone (Figure 1). Note that the model parameters of batch classifiers and cell-type/state classifiers are frozen during training, while only the GAN-based backbone parameters are updated in every epoch for ensuring batch-correction without losing the genuine image information (**Methods**). The batch distillation power of CytoMAD can be further enhanced by dividing the batch classifiers at the bottleneck region into multiple mini-classifiers and periodically retraining them at preset intervals (Figure 1c) (e.g., every 10 epochs (**Methods**)).

In summary, the self-supervised morphology distillator forms a dynamic feedback system with the GAN-based image translation to separate batch information from biological variations of interest, ultimately achieving batch-distilled phenotypic features and cell images. For the sake of clarity, we denoted the morphological profiles and cell images generated from the full model of CytoMAD as *CytoMAD-profiles* and *CytoMAD-images* respectively. We also assessed the performance of the classical GAN-based backbone without batch-aware morphology distillator (denoted as *no-CytoMAD*) for comparison, with its resultant phenotypic features and cellular images denoted as *no-CytoMAD-profiles* and *no-CytoMAD-images*.

### 2.2 Label-free Identifications of Human Lung Cancer Cell Types Across Data Sets

We first evaluated CytoMAD’s performance on classifying seven human lung cancer cell lines representing three major lung cancer types based on their label-free biophysical morphologies: lung squamous cell carcinoma (LUSC) (i.e., H520, H2170), adenocarcinoma (LUAD) (i.e., H358, H1975, HCC827), and small cell carcinoma (SCLC) (i.e., H69, H526). Three different image batches were captured (on different dates) for each cell line, deliberately introducing complex batch-to-batch variations arising from factors such as variations in the optical systems and microfluidic aspects. These diverse datasets are to test CytoMAD’s capability to identify the shared morphological profiles within the same lung cancer types, and thus faithfully distinguish different lung cancer types.

CytoMAD enables accurate image contrast translation (i.e., BF to QPI) and provides CytoMAD-images (i.e., CytoMAD QPI) (Figure 2a). Visually, CytoMAD QPI images closely resemble ground truth QPI. To quantify image contrast translation performance, we computed the structural similarity index (SSIM) and the root mean square error (RMSE) at the cell region (Figure 2b).

**Figure 2.**
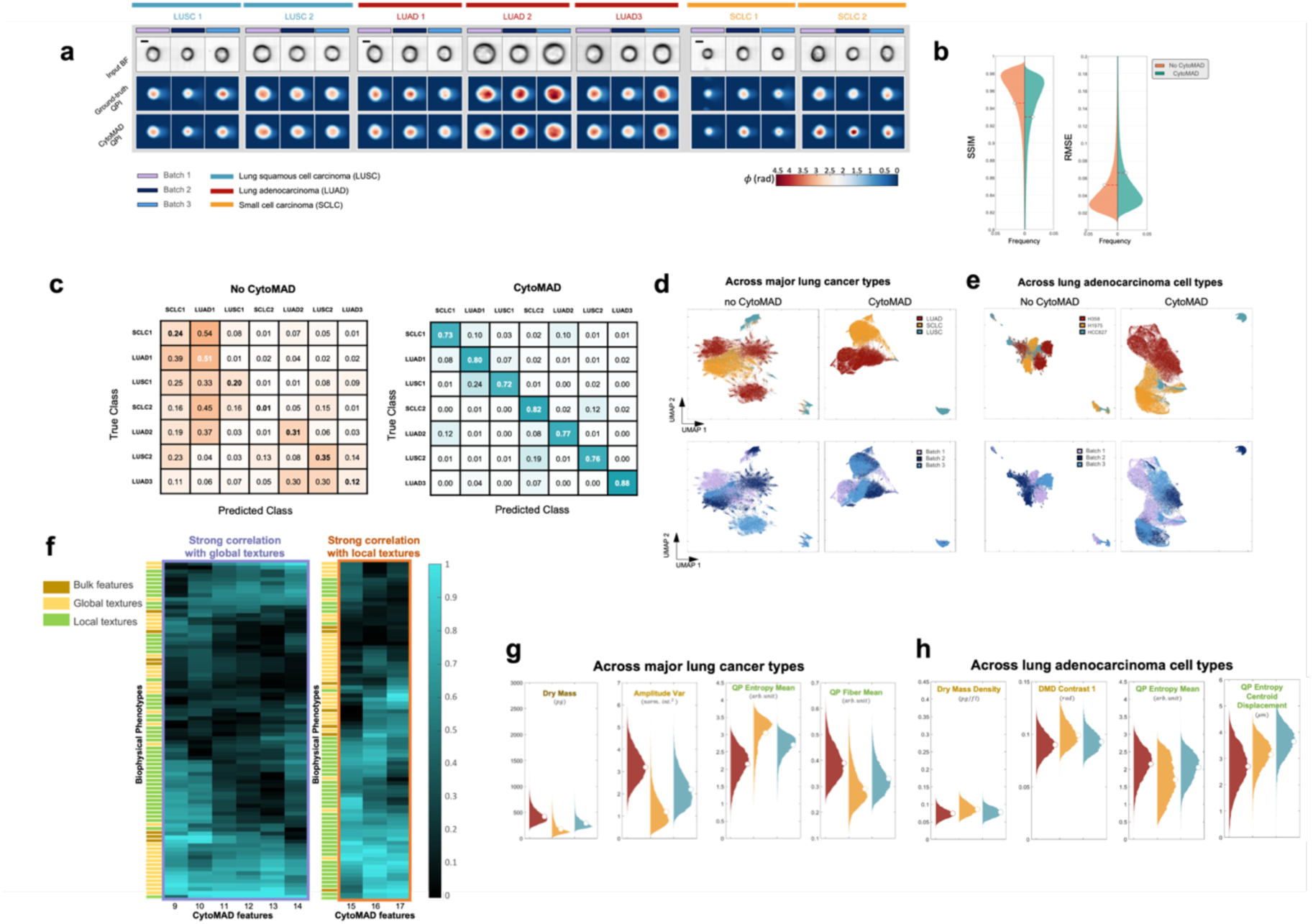
Batch Distillation Performance with CytoMAD on Lung Cancer Cell Lines Dataset. **(a)** Lung cancer cell lines images from multi-ATOM and CytoMAD results. The multi-ATOM label-free images of BF and QPI, and the CytoMAD-images (i.e., CytoMAD QPI) of the 7 types of lung cancer cell lines were reported. Color bar shows the phase values. Scale-bar: 10μm. FOV: 45μm × 45μm. **(b)** Violin plots on SSIM and RMSE distributions. The SSIM and RMSE distributions of no-CytoMAD-images are indicated in orange, while CytoMAD-images are indicated in green. The white circles represent the average values of the distributions. **(c)** Confusion matrix on lung cancer cell types classification with CytoMAD-profile. *Left:* Confusion matrix of no-CytoMAD-profile. *Right:* Confusion matrix with CytoMAD-profile. **(d)** UMAP across major lung cancer types. UMAP were computed with 63,000 cells data (i.e. 7,000 cells per cell type per batch). Every data point represents an individual cell. *Top-row*: Different colors are used to indicate different cancer types. *Bottom-row*: Different colors are used to indicate different batches. *Left*: UMAP of no-CytoMAD-profile. *Right*: UMAP with CytoMAD-profile. **(e)** UMAP across lung adenocarcinoma cell types. UMAP were computed with 63,000 cells data (i.e. 7,000 cells per cell type per batch). Every data point represents an individual cell. *Top-row*: Different colors are used to indicate different adenocarcinoma cell types. *Bottom-row*: Different colors are used to indicate different batches. *Left*: UMAP of no-CytoMAD-profile. *Right*: UMAP with CytoMAD-profile. **(f)** Selected absolute correlation profiles of between biophysical phenotypes and selected CytoMAD-profile group (See Supplementary Figure S2b for complete profile). The absolute correlation values are reported, with green color denotes a high correlation and black denotes a low correlation. Different colors are used to represent the 3 categories of the biophysical phenotypes. **(g)** Violin plots across major lung cancer types. **(h)** Violin plots across lung adenocarcinoma cell types.

We also compared the performance from the full model of CytoMAD and that of the classical GAN-based backbone without batch-aware morphology distillator (denoted as no-CytoMAD). The average SSIM for QPI of no-CytoMAD-images and CytoMAD-images was 0.9473 and 0.9305, respectively, indicating high structural similarity between the generated QPI and the ground truth QPI, and thus reliable image contrast conversion. The average RMSE values for QPI generated of no-CytoMAD-images and CytoMAD-images were 0.0519 and 0.0654, suggesting accurate phase value reconstructions. The similar SSIM and RMSE values in these two cases imply comparable reconstruction performance after adding the morphology distillation module in CytoMAD. The high SSIM and low RMSE values in CytoMAD QPI confirm the reliable image contrast translation from BF to QPI.

In the morphology distillator module, we trained cell type classifiers using CytoMAD-profile (or CytoMAD’s bottleneck latent features) (Figure 2c) and CytoMAD-images (**Supplementary Figure S1**) to assess biological information preservation while reducing batch-to-batch differences. Only one batch was used for training and validation, and classification performance was tested on two unseen batches (See **Methods**). For the cell type classifier based on CytoMAD-profile (Figure 2c), the across-batch classification accuracy was 0.2487 without CytoMAD (i.e., no-CytoMAD-profile), and it significantly improved to 0.7846 with CytoMAD, with all cell lines achieving >0.72 prediction accuracy. Classification performance based on CytoMAD-images was similar, with accuracies of 0.3280 without CytoMAD (i.e., no-CytoMAD-images) and 0.7768 with CytoMAD (**Supplementary Figure S1**). These substantial improvements in across-batch cell type classification, based on both CytoMAD features and images, demonstrate biological information preservation and reliability. Furthermore, this highlights CytoMAD’s batch-distillation power to overcome batch-to-batch variations.

We used Uniform Manifold Approximation and Projection (UMAP) for visualizing the mitigation of batch-to-batch variations and conservation of biologically relevant cell type information. UMAP analysis was conducted on the CytoMAD-profile across major lung cancer types (i.e., LUAD, LUSC, and SCLC) (Figure 2d) and different lung adenocarcinoma cell types (i.e., H358 (KRAS^G12C^), H1975 (EGFR^L858R/T790M^), HCC827 (EGFR^Del19^)) (Figure 2e). Without CytoMAD, we observed multiple clusters within the same type, especially LUAD (Figure 2d), indicating an obvious batch effect which obscures the difference among the three major cell type populations. In contrast, CytoMAD merges different batches of the same type into single clusters, and the three major cancer types became distinct. Likewise, in the LUAD cell type study (Figure 2e), without CytoMAD, multiple clusters within the same subtypes suggested strong batch differences. With CytoMAD, clusters unified, and the three subtypes appeared distinct. These findings reaffirm CytoMAD’s capability to minimize batch effects through batch-distilled phenotypic profiling.

We further investigate the interpretability of the self-supervised CytoMAD-profile in order to gain the model transparency and credibility, which are particularly pertinent in biomedical diagnosis. Specifically, we correlated the CytoMAD-profile with the hand-crafted biophysical phenotypes of cells from the CytoMAD output QPI (i.e., CytoMAD-images) and input BF images. These hand-crafted biophysical phenotypes of cells (a total of 84) features **(Supplementary Note S1)** were extracted based on a hierarchical morphological feature extraction approach [4, 7], which has recently shown promises in label-free single-cell morphological profiling. We identified the important CytoMAD-profile in classifying seven lung cancer cell lines based on feature importance and correlated them with hand-crafted biophysical phenotypes, which are categorized into 3 groups (i.e., bulk, global texture and local texture of biophysical morphology) (**Supplementary Figure S2a**). Specifically, hierarchical clustering grouped the selected CytoMAD-profile into five clusters (**Supplementary Figure S2b**). Notably, one group strongly correlated with global phenotypes, while another correlated with local biophysical phenotypes, offering insights into the biological relevance of CytoMAD-profile (Figure 2f).

We proceeded to analyze CytoMAD results for variations across major lung cancer types and LUAD cell types, as in the UMAP analysis. We compared biophysical phenotype distributions (readout from CytoMAD-images) across different cancer types (Figure 2g) and adenocarcinoma cell types (Figure 2h). Notable differences were observed in certain biophysical phenotypes across the three major cancer types (Figure 2g). They included the cell dry mass (bulk feature, effect size d = 0.40), opacity variance (global texture of cell opacity, effect size d = 0.57), and QP entropy mean and QP fiber mean (local textures of quantitative phase, or equivalently dry-mass density, effect size d = 0.44 and d = 0.30 respectively). In contrast, differences among LUAD cell types (Figure 2h) were more subtle, suggesting relatively similar cellular morphology across LUAD types compared to variations across major lung cancer types. (Figure 2h **& Supplementary Figure S3**).

### 2.3 Delineating Cellular Responses to Drug Treatments with CytoMAD

Morphological profiling is now becoming a promising technique in drug screening, yet batch effects pose a notable challenge. We assessed the CytoMAD approach using label-free drug response assays on LUSC (H2170) cells treated with docetaxel, afatinib, and gemcitabine at various concentrations across two batches, i.e., 5 concentration levels and 1 negative control) - forming 18 unique drug treatment conditions. Notably, during the CytoMAD model training, we provided only batch identifiers and drug types, purposely withholding drug concentration data. This approach was chosen to challenge the model’s capability for self-supervised learning of concentration-dependent morphological features (see **Supplementary Figure S4** for detailed training strategy).

In a comparative analysis of single-cell, label-free input images (BF and QPI) and CytoMAD QPI images (Figure 3a), we observed that CytoMAD QPI closely resembled the ground truth QPI, with a high SSIM of 0.9241 and low RMSE of 0.0098 (**Supplementary Figure S5**). These results reinforces the reliability of CytoMAD’s image contrast conversion in capturing contrast details pertinent to drug response assays.

**Figure 3.**
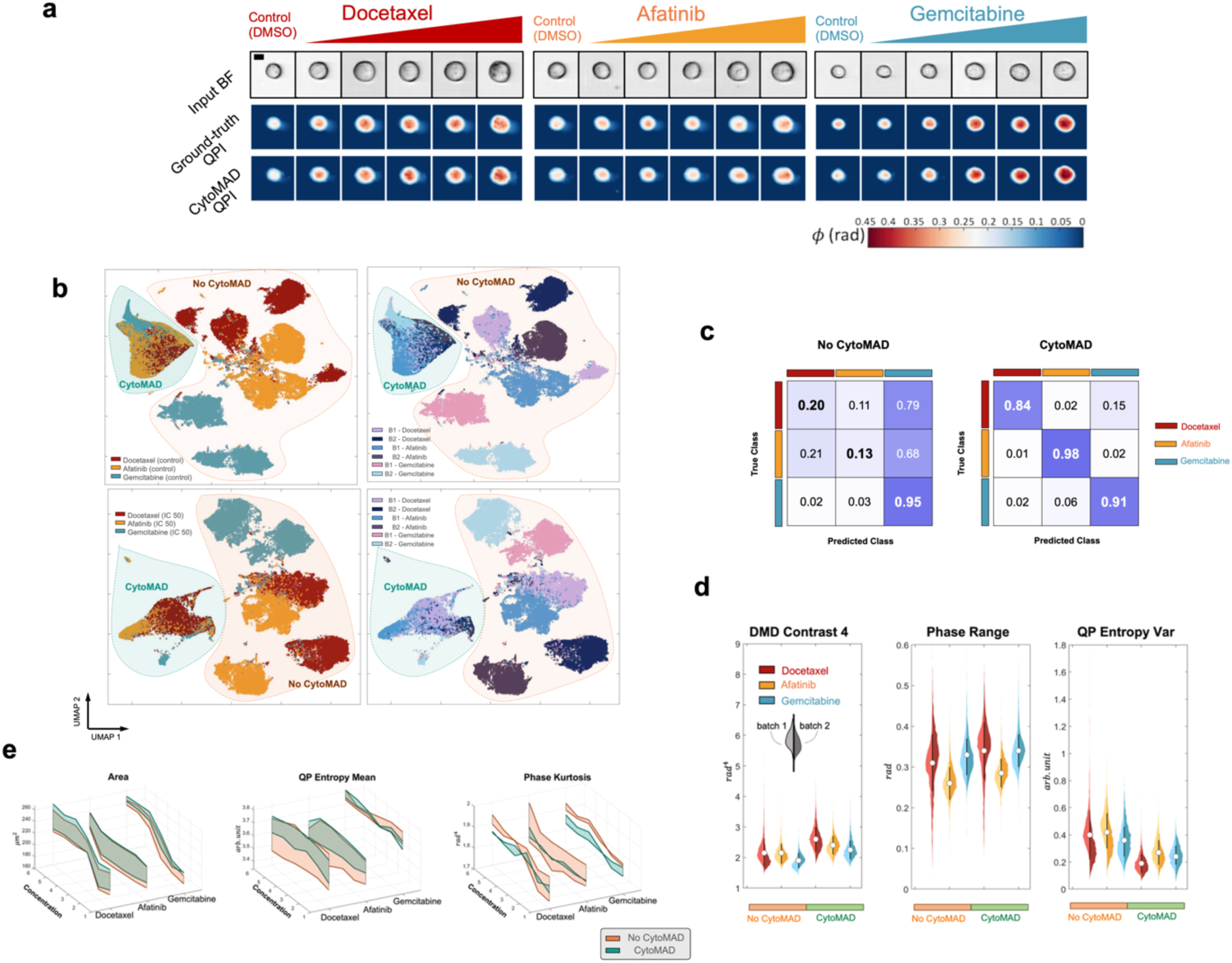
Drug Treatment Analyses with CytoMAD Batch Distillation. **(a)** H2170 drug response images from multi-ATOM and CytoMAD results. The multi-ATOM label-free images of BF and QPI, and the CytoMAD-images (i.e., CytoMAD QPI) of the H2170 drug treatment response were reported. Color bar shows the phase values. Scale-bar: 10μm. FOV: 45μm × 45μm. **(b)** UMAP across DMSO negative control samples and across IC50 samples. UMAP were computed with 60,000 cells data (i.e. 5,000 cells per batch without CytoMAD, and 5,000 cells per batch with CytoMAD). Every data point represents an individual cell. The orange circle denotes data without CytoMAD (i.e. no CytoMAD), while the green circle denotes data with CytoMAD. *Top-row*: UMAP across DMSO negative control samples. *Bottom-row*: UMAP across IC50 samples. *Left*: Different colors are used to indicate different drug treatment samples. *Right*: Different colors are used to indicate different batches. **(c)** Confusion matrix on IC50 samples with CytoMAD-profile. *Left:* Confusion matrix of no-CytoMAD-profile. *Right:* Confusion matrix with CytoMAD-profile. **(d)** Violin plots of biophysical phenotypes across IC50 samples. Different colors are used to indicate different drug treatment samples. The 2 sides of each violins represent 2 different batches and the white circle at the center line each violin denotes the average biophysical phenotype values between the 2 batches. The 3 violins on the left side of each plot indicates the biophysical phenotypic distribution without CytoMAD, while the 3 violins on the right side indicates the biophysical phenotypic distribution with CytoMAD. **(e)** Batch distance on biophysical phenotypes along drug concentration. Selected biophysical phenotypes, with their corresponding values in different drug treatments, were plot along the drug concentration. Each line on the plot represents a batch of each drug treatments and the shaded area between lines denotes the phenotypic distance between batches, with orange color denoting no CytoMAD and green denoting with CytoMAD.

To explore how well CytoMAD mitigates batch effects while maintaining critical cellular information, we performed UMAP analyses on negative control samples (DMSO) and IC50 samples of different drug treatments (Figure 3b). Prior to CytoMAD processing (i.e., no-CytoMAD) (orange area), samples from different batches appeared distinct on the UMAP plot, indicative of batch effects. After applying CytoMAD (green area), the variance attributed to batch effects was significantly reduced among control samples, evidenced by the convergence of samples from six batches into a unified cluster. Interestingly, when assessing the drug-treated IC50 samples, CytoMAD facilitated the formation of well-defined, drug-specific clusters (Green area in Figure 3b), accentuating its ability to discern treatment-specific morphological signatures while minimizing batch-related noise.

To further quantify the CytoMAD’s ability to delineate treatment responses, we next trained the drug treatment classifiers on IC50 samples. The training and validation sets consisted of a single batch with 2,000 and 500 cells per treatment, respectively (**Supplementary Note S5**). The across-batch classification, tested on an unseen batch with 2,500 cells per treatment, achieved accuracies of 0.91 with CytoMAD-profile, considerably higher than 0.43 with no-CytoMAD-profile (Figure 3c). This significant improvement underscores CytoMAD’s effectiveness in reducing batch differences while preserving treatment distinctions.

CytoMAD’s morphology-distillation ability was quantified by measuring differences in the hand-crafted biophysical phenotypes (extracted from CytoMAD images) among batches for each drug treatment (Figure 3d **& Supplementary Figure S6**). By quantifying the change in each biophysical phenotypic value after CytoMAD using a metric called, correction ratios (see **Methods**) (**Supplementary Figure S7**), we identified phenotypes with strong and weak batch corrections (Figure 3d & **Supplementary Figure S6**). Interestingly, we observed that the features related to high spatial-frequency information (e.g., DMD contrast 3, and QP entropy centroid displacement) exhibit higher correction ratio. It could indicate to that CytoMAD are trained to more effectively correct/reduce the high-frequency image noise in different batches.

Another evidence of CytoMAD’s effectiveness is the increased symmetry in violin plots after applying CytoMAD (comparing the left three violins with the right three) (Figure 3d). The CytoMAD plots exhibited a better symmetry when comparing the left and right violins, signifying a substantial reduction in inter-batch discrepancies, particularly for some biophysical phenotypes.

Based on our quantitative analysis that compared phenotypic distributions across drug treatments (represented by the violins of different colors in Figure 3d), the preserved patterns of the phenotypic distributions before and after CytoMAD show its ability to retain intrinsic cellular information for better discriminating the effect of different drug treatments. This is further corroborated by effect size measurements (after CytoMAD) for certain biophysical phenotypes across treatments include global dry-mass-density texture features (DMD contrast 4, effect size d = 0.19; phase range, effect, size d = 0.24) and local dry-mass-density texture features (QP entropy variance, effect size d = 0.17) (Figure 3d), which demonstrated considerable treatment-specific differences.

We further visualized how CytoMAD reduces the batch differences in different biophysical phenotypes along the drug concentrations (Figure 3e). By analyzing the trends of the biophysical phenotypes with different drug concentrations, we observe that CytoMAD effectively reduced the range of uncertainty in most of the biophysical phenotypes, as showed in the reduced shaded area (from orange to green) with CytoMAD, compared to that with no CytoMAD (Figure 3e **& Supplementary Figure S8).** It thus suggests CytoMAD’s capability to decrease batch differences in the biophysical phenotypes of cells, such as area (bulk features), phase kurtosis (global texture features), and QP entropy mean (local texture features). Importantly, similar trends along drug concentrations between the orange (without CytoMAD) and green (with CytoMAD) shaded areas demonstrate the preservation of progressive changes in the CytoMAD model, despite excluding the concentration information during training. All the above findings suggest insights into distinct label-free morphological responses to these drug treatments, consistent with their known different mechanisms of action (MoA).

### 2.4 Self-Supervised Label-free Biophysical Single-cell Analysis of NSCLC Biopsies

Investigating tumorigenesis in non-small-cell lung cancer (NSCLC), the leading cause of cancer-related mortality worldwide, is essential for unraveling the complex biological processes underlying tumor invasion, metastasis, and therapy resistance, all of which contribute to poor patient outcomes. Among these processes, epithelial-mesenchymal plasticity (EMP), particularly epithelial-mesenchymal transition (EMT), plays a pivotal role in driving tumor malignancy and invasiveness [28], characterized by the loss of epithelial markers and the gain of mesenchymal markers such as vimentin. In this study, we employ the CytoMAD method to investigate whether label-free biophysical cell morphologies can effectively capture subtle changes associated with EMP or related phenotypes in NSCLC biopsies from different patients.

Similar to the previous demonstrations, CytoMAD was trained to generate the batch-corrected morphological profiles (i.e., CytoMAD-profile) and predict the QPI (i.e., CytoMAD-images) from the input BF images. To better account for batch effects across highly heterogeneous clinical biopsies (four patient samples of lung adenocarcinoma obtained on four different dates in this study), we employed a self-supervised learning approach (Figure 4a). We first trained CytoMAD to perform a pretext task of classifying tumor biopsy versus blood samples from each patient.

**Figure 4.**
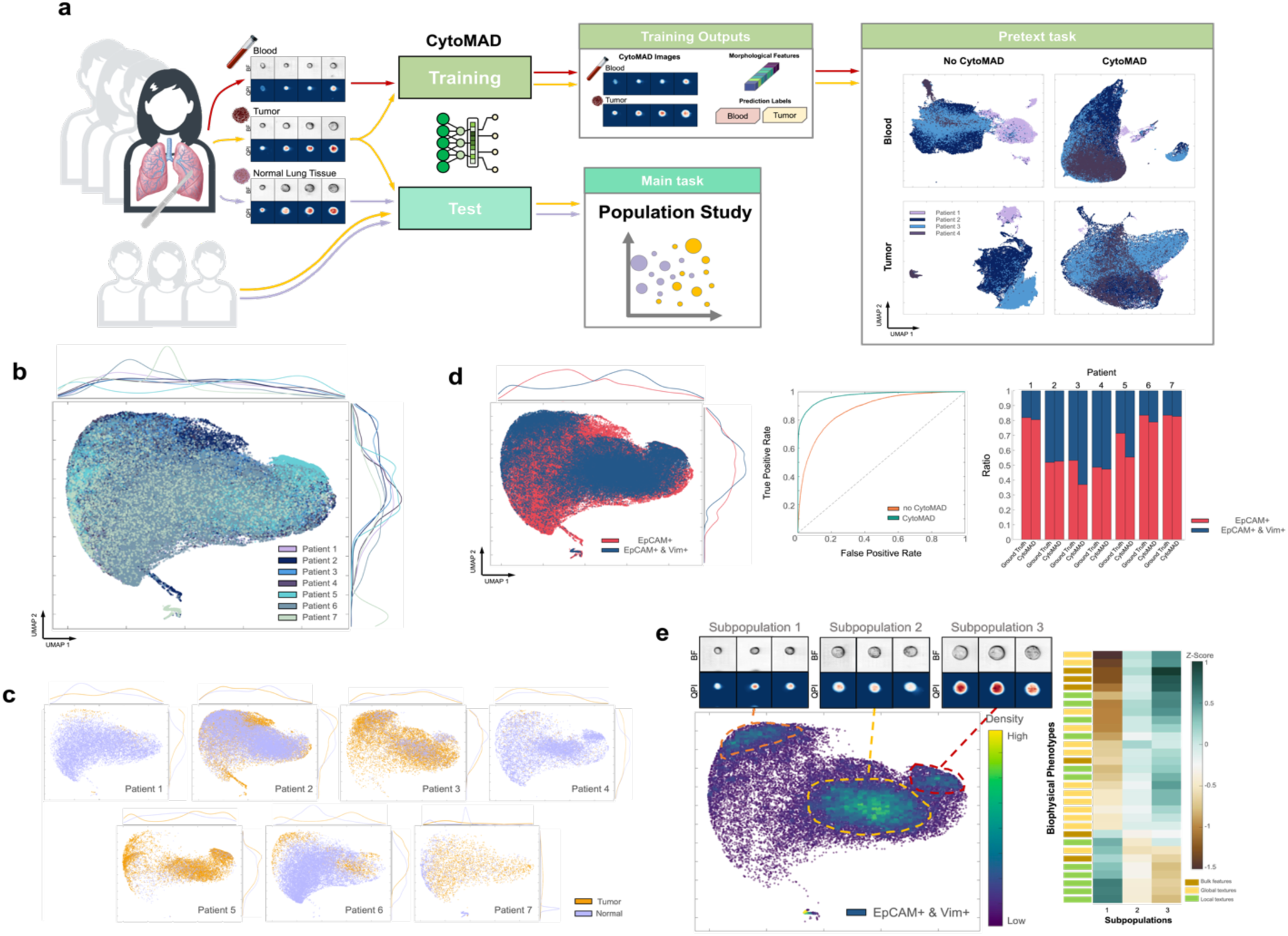
Pilot Study on NSCLC with CytoMAD. **(a)** NSCLC experimental pipeline with CytoMAD. With the collected samples (i.e. resected lung tumors tissue, normal lung tissue and peripheral blood samples) from 4 NSCLC patients, they were imaged with multi-ATOM for BF and QPI images. The tumor and blood data were employed in CytoMAD training to guide the disentanglement of valuable biological information from batch distortion during morphological distillation. Pretext tasks were conducted to validate the CytoMAD performance. Normal lung tissue samples, along with data from three additional NSCLC patients not included in the model training, were used to assess CytoMAD’s performance and test its generalizability across new patient populations. These samples were utilized for our main task, focusing on the population study involving both normal and tumor samples. **(b)** UMAP on tumor and normal lung tissue samples. Different colors are used to indicate different batches. **(c)** UMAP on tumor and normal lung tissue samples of each patient. Different colors are used to indicate different sample types. The distributions of each class are projected on the sides for visualization the highly-overlapping populations. **(d)** Molecular marker analysis of tumor and normal lung tissue samples. The UMAP, ROC and predicted population ratio of label-free molecular markers classification are reported. The true population ratio between EpCAM+ and Both+ cells is presented as a reference and different colors are used to denote the different molecular markers. **(e)** Subpopulation analysis in EMP cells. The UMAP of EMP cells (i.e. EpCAM+ & Vim+) are colored based on cell density. The BF and QPI of the subpopulations are reported. Scale-bar: 10μm. FOV: 45μm × 45μm. The z-score of the subpopulations are reported, with green color denotes a positive z-score and brown denotes a negative z-score. Different colors are used to represent the 3 categories of the biophysical phenotypes.

The rationale behind incorporating tumor biopsy and blood biopsy samples into CytoMAD training lies in the necessity for clear and distinct samples to guide the model in differentiating batch information from inherent cell characteristics. Acknowledging that tumor biopsy samples inherently contain a substantial number of non-tumor cells, and both tumor and normal lung tissue samples share common cell populations, our training strategy aimed to provide a more explicit ground truth for CytoMAD. Tumor cells and blood cells are generally very distinct in morphological characteristics, e.g., cell size, average QPI phase across the cell (**Supplementary Figure S9)**. They enabled us to work with samples exhibiting a marked distinction in cell types. Blood samples, easily accessible and featuring a highly heterogeneous mix of cell types, offered a clear contrast to tumor biopsy samples. This intentional selection of training samples not only aided the model in discerning batch information but also allowed CytoMAD to preserve more nuanced cell features. The inherent diversity in cell types within blood samples further contributed to the model’s effective generalization. Thus, the decision to employ tumor biopsy and blood biopsy samples as pretext task in CytoMAD training was guided by the need for unambiguous sample sets that could help the model accurately identify and minimize batch effects. The CytoMAD model was then tested to analyze the biophysical morphologies indicative of EMP-like phenotypes in both the resected tumor biopsies and the normal lung biopsies. We stress that the normal lung biopsy data was held out in training to assess the robustness of CytoMAD by analyzing unseen data.

First, the image translation performance (BF to QPI) proved to be robust across heterogeneous tumor biopsies, blood samples, and even unseen normal lung biopsies, achieving an average SSIM of 0.8881 and an RMSE of 0.0084 (**Supplementary Figure S10**). We further evaluated CytoMAD’s capability in correcting batch-to-batch variations among different patient batches based on the label-free CytoMAD-profile (Figure 4a**, and Supplementary Figure S11**). After applying CytoMAD, different batches, which were originally largely segregated (especially in tumor samples), exhibited improved merging, demonstrating the effectiveness of our approach in addressing batch effects.

Next, we investigated the CytoMAD-profile (i.e., the biophysical cell morphologies) of the tumor and normal lung biopsies. Using the fluorescence detection module integrated with the same imaging cytometry system (**Methods**), we first identified that significant cell populations overexpress both epithelial cell adhesion molecule (EpCAM+) and vimentin (Vim+), as well as EpCAM+ only (**Supplementary Figure S12**). Despite there are currently no specific molecular markers can universally define the mesenchymal state in all EMT programs [28], fluorescence labels targeting EpCAM and Vim are broadly regarded as valuable markers indicative of EMP - a partial EMT during which cells express with a mixture of epithelial and mesenchymal phenotypes [28]. With this knowledge, we sought to study the biophysical morphologies of the major populations of EpCAM+ and EpCAM+Vim+ cells in the normal and tumor biopsies, that has not been comprehensively investigated.

In the subsequent analysis, we added 3 new, unseen patients (denoted as patients 5, 6, and 7) to test the generalizability of the model across new patients, resulting in a total of 7 patient samples. These three additional patients were not part of the CytoMAD training process, serving as a real-life reference to test the model’s generalizability in downstream biological analysis. This scenario mimics the application of CytoMAD to new patient data without retraining the model, illustrating its effectiveness with previously unseen patients.

Based on the gating for EpCAM+ and EpCAM+Vim+ cells **(Supplementary Figure S12)**, we noted that various patient data batches exhibited reduced segregation in the UMAP embedding, showcasing the batch correction efficacy achieved by CytoMAD (Figure 4b & **Supplementary Figure S13)**. Notably, despite the exclusion of the three additional patients from the CytoMAD training set, their incorporation into the UMAP analysis also revealed a decreased level of batch segregation. This finding highlights CytoMAD’s capacity to effectively handle new, unseen patient data, showcasing its robust generalizability. The observed less-segregated clusters in the UMAP plot not only served as a validation but also emphasized CytoMAD’s efficacy in minimizing batch-to-batch variations across the patient datasets, even in the presence of new, previously unseen patient data. Interestingly, we could still discern some differences between tumor and the normal lung tissue populations across different patients (Figure 4c) - suggesting the ability of CytoMAD to explore the biophysical phenotypic differences between the normal and tumor tissue, echoing the recent work on label-free biophysical phenotyping of tumor biopsy [29].

Nevertheless, it should be recognized that tumor biopsies inherently heterogeneous, comprising of diverse of cell types (including non-cancerous cells), which could be attributable to the overlapping regions between tissue and normal tissue populations in the UMAP plot (Figure 4c). Recognizing this situation, our objective is to investigate the feasibility of using CytoMAD to identify the specific populations of cells with known molecular markers, i.e., EpCAM+ and the EpCAM+Vim+ cells. This could facilitate establishment of the biological grounding of label-free biophysical profile, extracted by CytoMAD, related to EMP (Figure 4d**, and Supplementary Figure S14).**

While the label-free CytoMAD-profile could generally reveal the segregation of the EpCAM+ and the EpCAM+Vim+ cells, we also observed a great degree of inter-patient variations **(Supplementary Figure S14)**. To delve deeper, we trained a deep neural network based on the label-free CytoMAD-profile to distinguish between EpCAM+ and EpCAM+Vim+ cells. The classification tests encompassed all 7 patients, ensuring a comprehensive evaluation of the model’s performance based on label-free CytoMAD-profile (Figure 4d). Using receiver operating characteristic curve (ROC) analysis, we observed a significant improvement in classification performance after CytoMAD, with an area under the curve (AUC) improvement from 0.8756 to 0.9705 (best at 1.00). Furthermore, the predicted population ratios between EpCAM+ and EpCAM+Vim+ cells based on the CytoMAD model resembled the ground truth in most of the patients (Figure 4d). These findings suggest that CytoMAD could faithfully preserve the biologically relevant information of biophysical cell morphologies across different clinical samples, i.e., reflecting the signature of EpCAM+ and EpCAM+Vim+ cells, and thus EMP-like phenotypes. They also substantiate the potential of using label-free biophysical cell morphologies to infer these molecular-specific cell phenotypes, distilled from the batch effect.

To explore further whether CytoMAD reveals additional biophysical insights that are otherwise absent in fluorescence markers. We examined the EpCAM+Vim+ cells and discovered three distinct subpopulations (Figure 4d-e **& Supplementary Figure S15**). Visually, subpopulation 1 predominantly consisted of smaller cells, while subpopulations 2 and 3 comprised progressively larger cells (Figure 4e). Furthermore, we observed distinct phenotypic differences between the three subpopulations based on the 84 biophysical features extracted from the CytoMAD output cell images **(Supplementary Note S1).** Based on the hierarchical feature categorization adopted earlier (Figure 2f), we found that the biophysical phenotypic differences between subpopulation 1 and subpopulations 2-3 (quantified by z-score) are mainly manifested by the bulk feature of Area and the global textures of dry mass density (e.g., Phase skewness, dry mass radial distribution). Conversely, subpopulations 2 and 3 exhibited similarities or less pronounced differences when compared to subpopulation 1. Noteworthy distinctions between subpopulations 2 and 3 primarily stemmed from cell orientation and cell fit texture, which captures the statistical moments of the profile that highlight the high spatial frequency of the phase. These findings demonstrate that the EpCAM+Vim+ cells, which might represent EMP, exhibit further heterogeneity in the biophysical traits that are not captured by the standard EpCAM and Vim markers. These findings thus underscore the power of label-free biophysical imaging, coupled with CytoMAD, in delineating cell heterogeneity and providing added values in analyzing tumor environments in the clinical samples. We note that, using the image-activated sorting modality recently developed [17, 30–38], any of these cell populations can be isolated according to the biophysical morphological features and analyzed for their molecular signatures (e.g., transcriptomes).

## 3. Discussion

In summary, CytoMAD introduces a new generative and integrative deep-learning approach that effectively tackles batch effects in image-based cytometry, while simultaneously facilitating image contrast translation to unveil additional cellular information and self-supervised morphological profiling. This work emphasizes its utility in the burgeoning field of biophysical cytometry, where CytoMAD augments BF image data to obtain quantitative biophysical phenotypes, such as cell mass, mass density, and their subcellular local and global distributions. This has profound implications in simplifying complex optical instrumentation required for conventional QPI operations, such as interferometric/holographic modules and multiplexed detection modules [26]. As a result, CytoMAD could pave the way for the broader adoption of biophysical cytometry across a wide range of applications, as evident by our demonstrations including accurate label-free classification of human lung cell types, functional drug-treatment assays, and biophysical cellular analysis of tumor biopsies from patients with early-stage NSCLC.

Batch effects have been extensively studied in other single-cell data modalities (e.g., single-cell omics), but remain largely uncharted in cell imaging, with few exceptions [13, 20]. In the current study, our primary emphasis was on addressing inter-batch variations, operating under the assumption that intra-batch variations are negligible. This assumption is supported by our comprehensive analysis of intra-batch variations in another comparable flow imaging cytometry technique (**Supplementary Figure S16**). CytoMAD’s unique ability to harness the power of self-supervised learning allows it to disentangle batch variations from biologically relevant morphological information during cross-modality image translation, enabling robust integrative image-based analysis across batches. We emphasize that CytoMAD is designed to mitigate rather than completely eradicate batch-to-batch variations. We showed that CytoMAD could still preserve subtle intra-batch heterogeneity (e.g., morphologically distinct populations of cells within the same batch (**Supplementary Figure S17**). This could ensure biological traits are retained while adeptly minimizes batch effects, enhancing data integration (e.g., the significant improvement in classification accuracy in Figure 2c **and 3c**) without indiscriminately conflating batches.

Without imposing any a priori assumptions on complex data distributions or requiring extensive manual annotation, CytoMAD delivers accurate QPI generation from BF images, as well as self-supervised batch-corrected morphological profiling for downstream analysis. All these aspects set CytoMAD apart from existing batch correction algorithms (**Supplementary Figure S18**). Our analysis showed that CytoMAD demonstrated superior performance in feature-based classification, achieving an across-batch accuracy of 0.91, surpassing HARMONY (0.78) [14] and MNN (0.70) [39] - the popular tools used in single-cell omics analysis (e.g., single-cell transcriptomics) (**Supplementary Figure S18**). We further note that these omics tools typically involve feature dimension reduction (through principal component analysis (PCA)) as the pre-processing step for stable batch correction. However, PCA collapses spatially structured data (from images) into linearly uncorrelated components that prioritize variance over biological significance. Hence, it could potentially overlook subtle yet important features and complicating the direct interpretation of variations to specific morphological traits. Moreover, these tools are not compatible with batch effect correction of 2D image data format. As a result, we stress that the comparisons shown in **Supplementary Figure S18**, while informative, should be regarded as indicative rather than definitive due to the absence of a direct, like-for-like benchmark.

Notably, CytoMAD can accurately predict progressive morphological changes in response to drug concentration trends, even without prior annotation (Figure 3). Utilizing blood and tumor classification as pretext tasks, CytoMAD effectively corrects batch effects and predicts label-free morphologies correlated with EpCAM and vimentin phenotypes in NSCLC biopsies (Figure 4).

Despite its promising results, CytoMAD could offer several opportunities for further development. Firstly, the cross-modality image translation/augmentation could be extended to other forms of contrast translation, particularly involving fluorescence image contrast, such as BF to fluorescence [21, 22, 24], QPI to fluorescence [23, 40, 41], or fluorescence to colorized BF [42, 43]. This augmentation approach could help establish a knowledge base that correlates and transfers molecular specificity into label-free morphological phenotypes of cells and tissues. Secondly, enhancing the interpretability of CytoMAD morphological profiles would enable more intuitive deciphering of underlying biological processes and mechanisms. Potential strategies to improve interpretability in CytoMAD include incorporating feature attribution methods (e.g., Layer-wise Relevance Propagation (LRP) [44], Gradient-weighted Class Activation Mapping (Grad-CAM) [45]) to visualize influential regions within input images, providing insights into learned representations; or integrating disentangled learning techniques within the autoencoder architecture to learn more interpretable and independent features that might be more readily linked to underlying biology.

Overall, our findings demonstrate that CytoMAD heralds a new development of deep-learning-based, batch-aware morphological profiling of cells. Its application in biophysical cytometry highlights its potential for accurate and insightful biophysical investigations into complex biological processes. It might enrich our understanding of cellular functions and inspire the discovery of cost-effective biomarkers for diagnostic and therapeutic purposes.

## 4. Methods

### 4.1 Overall Framework of CytoMAD

CytoMAD is a generative deep-learning model tailored for batch-aware cell morphology distillation. It is built upon a conditional GAN (cGAN) of a Pix2Pix architecture [46] integrated with a set of classification networks, altogether enabling robust image contrast translation and distillation of biological phenotypes from batch variations. CytoMAD offers both batch-distilled phenotypic features (i.e., CytoMAD-profile) and cellular images (i.e., CytoMAD-images) as model output. These results could further be utilized for downstream biological analysis. In general, the CytoMAD model comprises two synergistic functions: (i) the GAN-based backbone for image generation and contrast translation, and (ii) the classifier-guided, batch-aware morphology distillation (Figure 1b).

#### 4.1.1 Pre-training CytoMAD for Image Contrast Translation

With cGAN as the backbone, the CytoMAD model comprises a generator network for image-to-image translation and a discriminator classifier for optimizing the generator prediction through a feedback mechanism. Before implementing the batch-aware module, the model is initially pre-trained for image generation and contrast translation.

The generator takes cell images captured in a particular image contrasts (e.g., brightfield (BF)) as the model input. The images are then directed to the encoder and undergo multiple layers of 2D convolutional layers, batch normalization layers, and mathematical activation functions. These layers condense the biological information from the input images into a 1D array at the bottleneck layer. Such an array encodes the important features representing the input cell images and, thus, serves as the cellular phenotypic features, which is referred to no-CytoMAD-profile at this pre-training stage. The output images at this pre-training stage, called no-CytoMAD-images, are reconstructed based on this 1D information with an additional image contrast translation step (i.e., from BF to quantitative phase image (QPI) in this paper). The array passes through multiple deconvolutional layers, batch normalization layers, and mathematical activation equations at the decoder for image reconstruction and translation. Skip-in layers are implemented between the encoder and decoder for better image feature preservation **(Supplementary Note S2)**.

For the discriminator network, it improves the image reconstruction/translation of the generator by distinguishing its predicted images from original target images (i.e., original “ground-truth” QPI (Figure 1)), hence, forming a feedback mechanism with the classification loss for optimizing the generator parameters. Hence, the CytoMAD model can be pre-trained for accurate image reconstruction and contrast conversion from BF to QPI. The contrast translation serves as an additional feature of the CytoMAD for augmented cellular information, in conjunction with its batch-aware capability. In the applications where image contrast conversion is not required, convolutional autoencoder architecture could be adopted by equalizing the input and output target images of the cGAN backbone.

#### 4.1.2 Classifier-guided Batch-aware Morphology Distillation

CytoMAD differs from the existing GAN models by the implementation of morphology distillator. The morphology distillator is a set of classification networks, which learn in synergy to minimize batch-to-batch variations while preserving biological information in the images. These classification networks take a crucial role here through 2 types of classifiers: batch classifiers and cell-type/state classifiers (Figure 1).

These classifiers are first pre-trained based on no-CytoMAD-profile and no-CytoMAD-images to identify the batch and cell-type information in the cGAN backbone. They are then implemented at both the bottleneck region and the output of the generator model, with the *frozen* models’ parameters, for guiding the next training round of batch correction in the generator model. By freezing the classifiers’ parameters and including the morphology distillatory loss into the CytoMAD loss function, we enforce the generator to recognize and remove batch information while preserving cell-type information with minimal alterations with the no-CytoMAD-profile and no-CytoMAD-images.

At the bottleneck region of the pre-trained generator, the batch classifiers aim to reduce the batch-to-batch variations, while the cell-type/state classifiers learn to preserve the cellular variations in a 1D feature format, which is referred to the CytoMAD-profile. These classifiers follow the basic framework of neural networks, which is detailed in **Supplementary Note S3**. Since the output images are reconstructed chiefly based on the bottleneck 1D features, these classifiers promote the disentanglement between the batch information and cell morphological information. Hence, they facilitate the morphology-distillation process, uncovering valuable morphological information in both the 1D CytoMAD profile and 2D batch-aware cell images (i.e., CytoMAD-images).

Additional batch and cell-type/state classifiers based on convolutional neural networks are also present at the end of the generator **(Supplementary Note S3)**. They guide the reconstruction of batch-aware cell images (CytoMAD-images) and remove the batch information induced by the encoder-decoder skip-in layers. Employing multiple batch classifiers and periodically retraining them at predetermined intervals (e.g., every 10 epochs) in the CytoMAD model can facilitate and expedite the batch correction procedure.

During the CytoMAD training, the model parameters of batch classifiers and cell-type/state classifiers within the morphology distillator are frozen, while only the generator’s parameters and the discriminator’s parameters are updated in every epoch for batch-correction and ensuring the image prediction accuracy.

Overall, all these classification networks contribute to the CytoMAD loss function *L_cytoMAD_*

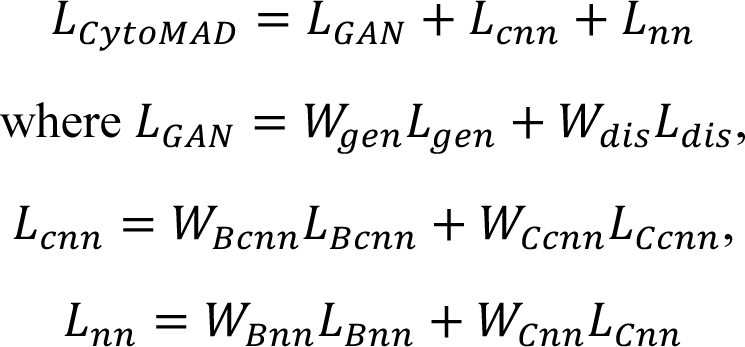

*L_GAN_* is the loss of GAN-backbone model, with *W_gen_* and *L_gen_* denote the weighting and the mean square error loss of generator model respectively, *W_dis_* and *L_dis_* denote the weighting and the binary cross entropy loss of discriminator model. *L_cnn_* is the loss of convolutional-neural-network-based classifier models, with *W_Bcnn_* and *L_Bcnn_* denote the weighting and the categorical cross entropy loss of batch classifier model, *W_Ccnn_* and *L_Ccnn_* denote the weighting and the categorical cross entropy loss of cell type classifier model. *L_nn_* is the loss of neural-network-based classifier models, with *W_Bnn_* and *L_Bnn_* denote the weighting and the categorical cross entropy loss of batch classifier model, *W_Cnn_* and *L_Ccnn_* denote the weighting and the categorical cross entropy loss of cell type classifier model. All these networks form an integrated dynamic feedback system with the GAN-based image translation for extracting the batch information from the biological variations of interest and, eventually, achieving batch-aware capability for morphological CytoMAD profiling and the image contrast translation.

### 4.2 Assessment of CytoMAD Performance: Image Contrast Translation Accuracy

We employed two metrics to evaluate the performance image translation (BF to QPI) by quantifying the similarity and difference in image pixel values between the original images and the CytoMAD images (i.e. the phase values).

#### 4.2.1 Structural Similarity Index Measure (SSIM)

SSIM is a perceptual metric broadly adopted for quantifying similarities within pixels between images [47]. It evaluates how well the image structure in CytoMAD images is preserved from original target images (i.e. QPI). Given that the valuable biological information resides in the cell region and the downstream analysis is conducted based on this region, the SSIM values were calculated and reported based on the cell area only through cell segmentation to differentiate cell bodies from the background. A high SSIM value of approaching 1 represents a high similarity between images.

#### 4.2.2 Root Mean Square Error (RMSE)

RMSE was also used for computing the pixel to pixel values difference between the original images and the CytoMAD images. Similarly, RMSE were reported based on the cell region only for comprehensive study.

### 4.3 Assessment of CytoMAD Performance: Morphology Distillator

We examined the batch effect removal efficiency of CytoMAD through the visualization given by Uniform Manifold Approximation and Projection (UMAP) and quantification of biophysical phenotypic profile correction.

#### 4.3.1 UMAP analysis of batch-effect reduction and biological information preservation

To evaluate the ability to mitigate the batch-to-batch variations and to simultaneously preserve the biological information, UMAP analyses were conducted based on the no-CytoMAD-profile and CytoMAD-profile. This visualization enables the observation of the degree of batch mixing across the multiple batches, and hence, visualizing the batch effect removal efficiency of CytoMAD. At the same time, it also allows direct visualization of the how well the data of the same cell types/state across the multiple batches could cluster together in UMAP - an indication of the ability to preserve the biologically relevant information in the data.

#### 4.3.2 Quantification of batch-effect reduction in the biophysical profiles

In addition to UMAP examination, we also conducted quantitative analysis to assess the batch effect reduction efficiency. This was achieved by first extracting the biophysical morphological profiles from both the CytoMAD-images and the original QPI. This profile parametrized a catalog of 84 biophysical features of cells following a spatial hierarchical manner, including bulk phenotypes (e.g. area, circularity), global phenotypes (e.g. dry mass density, attenuation density) and local phenotypes (e.g. BF entropy, phase entropy) [4, 7]. We thus could quantify the mean values of each biophysical feature within each batch of samples. We then defined “batch distance” as the difference in mean values across batches of the same samples. We further compared the batch distance of each biophysical features between the original QPI and the CytoMAD images. The reduction in batch distance thus indicates the decrease of batch-to-batch variations across samples.

Here the rationale of utilizing biophysical profile instead of bottleneck CytoMAD profile for assessing the batch-effect reduction performance is that biophysical profiles are constructed based on the well-defined geometrical metrics as well as human interpretable features. This makes them well-suited for drawing biological interpretation in the detailed morphological analysis (e.g. Figure 2f-h, Figure 3d-e, Figure 4d). By computing the batch distance reduction based on biophysical profile from CytoMAD images, we can gain further insights into which specific biophysical phenotypes are more prone to batch effect and demonstrate stronger batch correction with CytoMAD. This analysis offers valuable cues on the impact of batch effect on different biophysical profile and the efficacy of CytoMAD in overcoming these differences on each biophysical feature (**Supplementary Figure S6-8**).

#### 4.3.3 Across-batch Cell Type/State Classification

The capability of biological information preservation was evaluated through cell type/state classification using both the CytoMAD profiles and CytoMAD images. Deep neural networks were used for the classification based on CytoMAD profiles. The model consists of 3 dense layers with 75, 50 and 25 nodes, interconnected with rectified linear unit (ReLU) activation function. As for CytoMAD image classification, we employed the convolution neural network, which is a 5 layers model for image-based classification, with each of the layers composed of 2D convolution, batch normalization, leaky ReLU activation functions and max pooling operation. Both the deep neural network models and the convolution neural network models were trained for 100 epochs, utilizing softmax function as the output activation function and categorical cross-entropy loss as the loss function.

To better assess the preservation of biological information while reducing the batch-to-batch differences, the cell type/state classifiers were trained with one batch/selected batches of samples only and tested using unseen batches to evaluate the across-batch classification performance.

### 4.4 Multi-ATOM imaging

Multi-ATOM combines the time-stretch imaging technique [48, 49] and phase gradient multiplexing method to retrieve complex optical field information (including the BF and quantitative-phase contrasts) of the cells at high speed in an interferometry-free manner (Figure 1a). Detailed working principle and experimental configuration were reported previously [2, 48]. In brief, a wavelength-swept laser source was firstly generated by a home-built all-normal dispersion (ANDi) laser (centered wavelength: 1064 nm; bandwidth: ∼10 nm; repetition rate: 11 MHz; pulse width = ∼12 ps). The laser pulses were temporally stretched in a single-mode dispersive fiber, and were then amplified by an ytterbium-doped fiber amplifier module. The pulsed beam was subsequently launched to and spatially dispersed by a diffraction grating into a 1D line-scan beam which was projected orthogonally onto the cells flowing in the customized microfluidic channel. This line-scan beam was transformed back to a single collimated beam after passing through a double-pass configuration formed by a pair of objective lenses (N.A. = 0.75/0.8). Afterwards, the beam conveying phase-gradient information of the cell was split into 4 replicas by a one-to-four de-multiplexer, where each beam profile was half-blocked by a knife edge from 4 different orientations (left, right, top and bottom) respectively. Recombining the 4 beams by a four-to-one fiber-based time-multiplexer, we were able to detect the line-scan phase-gradient information in 4 directions in time sequence at high speed by a single-pixel photodetector (electrical bandwidth = 12 GHz). The digitized data stream was processed by a real-time field programmable gate array (FPGA) based signal processing system (electrical bandwidth = 2 GHz, sampling rate = 4 GSa/s) for primary cell detection and image segmentation with a processing throughput of >10,000 cells/s in real-time. These segmented phase-gradient images of cells were sent to four data storage nodes (memory capacity > 800 GB) through four 10G Ethernet links, which were reconstructed to 2D complex field information following a complex Fourier integration algorithm, detailed elsewhere [27]. The multi-ATOM system also includes a fluorescence detection module, similar to the previous work [4]. In brief, two continuous wave (CW) lasers (wavelength: 488nm and 532nm) were employed to generate line-shaped fluorescence excitation, that were spatially overlapped with the multi-ATOM illumination. The two epi-fluorescence signals were detected by two photomultiplier tubes (PMT) separately. In the analog electronics backend, we multiplexed the PMT-detected signals by frequency modulation (11.8 MHz and 35.4 MHz respectively, using a multichannel direct digital synthesizer). The multiplexed signals were then separated by digital demodulation and low-pass filtering. The same FPGA was configured to synchronously obtain the signal from multi-ATOM and fluorescence detection from each single cell at high-speed.

### 4.5 Microfluidic chip fabrication

The detailed microfluidic chip fabrication steps can be referred to our previous work [4]. In brief, the microfluidic channel was fabricated using standard soft lithography on a silicon wafer mold. A layer of photoresist was applied to a silicon wafer using a spin coater, followed by a two-step soft-bake. The photoresist was then patterned with a maskless soft lithography machine. The exposed photoresist was post-baked and developed. Subsequently, a polydimethylsiloxane (PDMS) precursor was applied to the silicon wafer and cured to form the channel. Post-curing, the channel was demolded and prepped for plastic tubing insertion. The channel was bonded to a glass slide using oxygen plasma, followed by an oven bake to strengthen the bonding. The microfluidic chip contains a serpentine channel structure, critical for robust in-focus single-cell imaging, which included 8 repeated units. The channel dimension at the imaging section was 30 mm x 60 mm.

### 4.6 Datasets

Effective training of deep learning models generally necessitates large datasets. In this regard, the high-throughput nature of our multi-ATOM imaging flow cytometry plays a vital role in generating large-scale, label-free cell images at ultrahigh-throughputs of >10,000 cells/sec. Prior to CytoMAD training, these large-scale single-cell image datasets (captured from in-vitro cultured cells, as well as the clinical patient samples) were first screened by a pre-processing pipeline for data quality control (**Supplementary Figure S19**). Cell segmentation was first applied to the BF and QPI images captured by multi-ATOM to generate the cell body masks, which facilitated the definition of biophysical phenotypes as well as the a set of focusing factors. A total 84 biophysical phenotypes, which represent the cell morphological properties, were defined and grouped into 3 hierarchical categories: bulk phenotypes (e.g. area, circularity), global phenotypes (e.g. dry mass density, attenuation density) and local phenotypes (e.g. BF entropy, phase entropy). Furthermore, we also defined a set of cell focusing factors from the segmented masks that quantify the image focusing quality and thus to help exclude out-of-focused cell image. These focusing factors, considering various aspects such as gradient-based measures and intensity statistics within the cell regions, aim to assess the sharpness and clarity of cellular features. By incorporating both spatial gradients and pixel intensities, these factors provide a comprehensive quantitative evaluation of image focus quality. All these parameters were then used to generate 2-dimensional scatter plots for preliminary visual image inspection and and cell gating to exclude cells of out-of-focused or debris. Subsequently, all the remaining cells within the population were selected for the CytoMAD training/testing and the downstream analysis. This pre-processing pipeline ensured data quality across all multi-ATOM imaging datasets, laying a robust foundation for quantitative cell analysis.

To produce multiple batches of data for the CytoMAD’s demonstration, cell images were collected on different dates using multi-ATOM. This approach intentionally introduced inherent, intricate batch-to-batch variations attributed from various sources, including (1) the optical system factors such as power instability of the laser source (e.g., the intensity noise of the pulsed laser) and the noise originated from photodetection and signal amplification in the system; (2) the microfluidic aspects associated with the variations in the microfluidic chip fabrication quality that could potentially result in the image distortion/aberration. This deliberately introduced complex batch-to-batch variations, aiming to validate CytoMAD’s ability in minimizing batch effects and provide a realistic scenario regarding integrated analysis across datasets.

#### 4.6.1 Lung Cancer Cell Lines

In this study, a total of 7 lung cancer cell lines were imaged by multi-ATOM and analyzed on 7 different days, producing 3 batches of ∼120,000 cells per cell line (i.e. >1,000,000 single-cell images in total, each of which consists of two label-free contrasts: BF and QPI). These cell lines represent three major lung cancer types: lung squamous cell carcinoma (LUSC) (i.e., H520, H2170), adenocarcinoma (LUAD) (i.e., H358, H1975, HCC827), and small cell carcinoma (SCLC) (i.e., H69, H526). The CytoMAD model was trained with 1,000 cells per cell line per batch, validated with 200 cells per cell line per batch and tested with ∼40,000 cells per cell line per batch (see **Supplementary Note S5)**.

#### 4.6.2 Drug assays of H2170

In this experiment, H2170 were treated with three drugs, which have different mechanism of action (MoA), i.e. Docetaxel as microtubule stabilizing agent [50], Afatinib as tyrosine kinase inhibitor in targeted therapy [51] and Gemcitabine as antimetabolite [52]), each with 5 concentration levels and a negative control with dimethyl sulfoxide (DMSO) treated for 24 hours (see **Supplementary Note S6)**. They were imaged using multi-ATOM for single-cell BF and QPI images on 6 days, forming 2 batches with ∼100,000 cells per drug. Basically, this dataset consists of 2 batches of data, with each batch containing 3 different drug treatments and each treatment comprising 6 different concentration conditions. This resulted in 18 unique drug treatment conditions in each batch.

In contrast to the previous lung cancer sub-type classification, this drug assay study focused on the ability of CytoMAD to reveal subtle and progressive changes in response to different drug concentration. The CytoMAD model was trained with 1,000 cells per drug treatment conditions per batch, validated with 500 cells per cell line per drug treatment conditions per batch, and tested with 5,000 cells per drug treatment conditions per batch (see **Supplementary Note S5)**. To examine the capability of CytoMAD in preserving progressive changes along drug concentration, only the batch information and the drug treatment types (i.e. docetaxel, afatinib, gemcitabine, and control) were provided to the model for guiding the batch-aware morphology distillation. The drug concentration information was held out to assess the power of CytoMAD.

#### 4.6.3 Lung Cancer Patients Samples

We recruited and collected samples from 7 NSCLC patients diagnosed with adenocarcinoma in the Queen Mary Hospital of Hong Kong, with each of them composed of resected lung tumors tissue, normal lung tissue and 9 mL of peripheral blood samples. They were then imaged using multi-ATOM on separated date, resulting in 4 batches of data with a total of ∼180,000 cells. Written consents for clinical care and research purposes were obtained from the donors. The research protocol was approved by the University of Hong Kong/Hospital Authority Hong Kong West Cluster Institutional Review Board (UW 19-278). After the standard sample pre-processing (e.g. disaggregating tissue into single-cell suspension, red blood cell lysis), the patient samples were imaged with multi-ATOM on separated dates, forming 7 batches with 2 label-free contrast images of BF and QPI of ∼180,000 cells. For each patient, their samples (i.e., tumor sample, normal lung tissue sample and peripheral blood sample) were collected on the same days and under the same optical system settings. Thus, negligible batch effect across these samples from the same patient.

CytoMAD model was trained based on tumors and blood samples from 4 patients, comprising 1,000 cells and 500 cells per sample per patient for training and validation respectively. Its performance was then assessed with >120,000 cells from resected tumors and peripheral blood, in addition to ∼56,000 cells from unseen (held-out) patients’ normal lung tissue (see Supplementary Note S5).

### Data availability

Sample data of 7 lung cancer cell lines dataset and the drug treatment response of H2170 dataset are available at https://github.com/MichelleLCK/CytoMAD. The full datasets will be released upon request.

### Code availability

CytoMAD code and sample data are available at https://github.com/MichelleLCK/CytoMAD with guidelines and tutorials.

## Acknowledgement

The work is supported by the Research Grants Council of the Hong Kong Special Administrative Region of China (grant nos. 17125121, RFS2021-7S06, C7047-16G), Platform Technology Funding of the University of Hong Kong.

## Author Contributions

K.K.T. conceived the project. M.C.K.L. M.C.K.L. and D.M.D.S. designed and performed experiments, with advices from K.K.T. M.C.K.L. performed data analysis with interpretation from D.M.D.S., M.K.Y.H., and J.C.M.H. M.C.K.L. and K.K.T. wrote the manuscript, with assistance from D.M.D.S., K.C.M.L., J.S.J.W., M.K.Y.H., and J.C.M.H. All authors edited the manuscript.

