## Supplementary Information for "Information-Distilled Generative Label-Free Morphological Profiling Encodes Cellular Heterogeneity"

#### Table of Content

**Supplementary Note S1:** Full list of biophysical phenotypes and their hierarchical categories

**Supplementary Note S2:** Model architecture of CytoMAD generator

**Supplementary Note S3:** Model architecture of classifiers in CytoMAD morphology distillator

**Supplementary Note S4:** Training details and hyperparameters of CytoMAD

**Supplementary Note S5:** Training data size of CytoMAD

**Supplementary Note S6:** Concentration Conditions of Drug Assays of H2170

**Figure S1:** Confusion matrix on lung cancer cell types classification with CytoMAD QPI

**Figure S2:** Correlation between CytoMAD-features profile and biophysical phenotypes

**Figure S3:** CytoMAD-based biophysical phenotypes across major lung cancer types and adenocarcinoma subtypes

**Figure S4:** H2170 drug assays experimental design with CytoMAD

**Figure S5:** SSIM and RMSE distributions of H2170 in the drug assays

**Figure S6:** Violin plots of biophysical phenotypes across IC50 samples

**Figure S7:** Batch distance correction ratio after CytoMAD

**Figure S8:** Batch distance on biophysical phenotypes along drug concentration

**Figure S9:** NSCLC samples images from multi-ATOM and CytoMAD-images results

**Figure S10:** SSIM and RMSE of NSCLC samples

**Figure S11:** UMAP on NSCLC blood and tumor samples

**Figure S12:** NSCLC EpCAM and vimentin fluorescence signals and gating

**Figure S13:** UMAP on different batches/patients of tumor and normal lung tissue samples

**Figure S14:** UMAP on molecular markers of tumor and normal lung tissue samples

**Figure S15:** Z-Score of subpopulation analysis in EpCAM+Vim+ cells

**Figure S16:** UMAP on intra-batch analysis

**Figure S17:** UMAP on H520 cell samples with CytoMAD

**Figure S18:** Benchmarking with existing batch-correction algorithms

**Figure S19:** Pre-processing pipeline of multi-ATOM images

#### Supplementary Note S1: Full list of biophysical phenotypes and their hierarchical categories

| | | |
|---|---|---|
| **Bulk** | | Area |
| | | Volume |
| | | Circularity |
| | | Eccentricity |
| | | Aspect Ratio |
| | | Orientation |
| | | Dry Mass |
| **Global** | **Optical Density** | Attenuation Density |
| | | Amplitude Var |
| | | Amplitude Skewness |
| | | Amplitude Kurtosis |
| | | Peak Amplitude |
| | | Peak Absorption |
| | | Amplitude Range |
| | **Mass Density** | Dry Mass Density |
| | | Dry Mass Var |
| | | Dry Mass Skewness |
| | | Dry Mass Radial Distribution |
| | | Dry Mass Centroid Displacement |
| | | Peak Phase |
| | | Phase Var |
| | | Phase Skewness |
| | | Phase Kurtosis |
| | | Phase Range |
| | | Phase Min |
| | | Phase Radial Distribution |
| | | Phase Centroid Displacement |
| | | Mean Phase Arrangement |
| | | Phase Arrangement Var |
| | | Phase Arrangement Skewness |
| | | Phase Orientation Var |
| | | Phase Orientation Kurtosis |
| **Local** | **Optical Density** | BF Entropy Mean |
| | | BF Entropy Var |
| | | BF Entropy Skewness |
| | | BF Entropy Kurtosis |
| | | BF Entropy Range |
| | | BF Entropy Peak |
| | | BF Entropy Min |
| | | BF Entropy Centroid Displacement |
| | | BF Entropy Radial Distribution |
| | | BF STD Mean |
| **Local** | **Optical Density** | BF STD Var |
| | | BF STD Skewness |
| | | BF STD Kurtosis |
| | | BF STD Range |
| | | BF STD Peak |
| | | BF STD Min |
| | | BF STD Centroid Displacement |
| | | BF STD Radial Distribution |
| | | BF Fiber Texture Centroid Displacement |
| | | BF Fiber Texture Radial Distribution |
| | | BF Fiber Texture Pixel >Upper Percentile |
| | | BF Fiber Texture Pixel >Median |
| | | BF Fiber Mean |
| | | BF Fiber Variance |
| | | BF Fiber Skewness |
| | | BF Fiber Kurtosis |
| | **Mass Density** | Phase STD Mean |
| | | Phase STD Var |
| | | Phase STD Skewness |
| | | Phase STD Kurtosis |
| | | Phase STD Centroid Displacement |
| | | Phase STD Radial Distribution |
| | | Fit Texture Mean |
| | | Fit Texture Variance |
| | | Fit Texture Skewness |
| | | Fit Texture Kurtosis |
| | | Fit Texture Centroid Displacement |
| | | Fit Texture Radial Distribution |
| | | Phase Entropy Mean |
| | | Phase Entropy Var |
| | | Phase Entropy Skewness |
| | | Phase Entropy Kurtosis |
| | | Phase Entropy Centroid Displacement |
| | | Phase Entropy Radial Distribution |
| | | Phase Fiber Centroid Displacement |
| | | Phase Fiber Radial Distribution |
| | | Phase Fiber Pixel >Upper Percentile |
| | | Phase Fiber Pixel >Median |
| | | Phase Fiber Mean |
| | | Phase Fiber Var |
| | | Phase Fiber Skewness |
| | | Phase Fiber Kurtosis |

We defined a comprehensive set of biophysical phenotypes to delineate the cell morphological and biological properties based on single-cell brightfield (BF) and quantitative phase images (QPI) from multi-ATOM system. Within these 84 biophysical features, they could be grouped into 3 hierarchical categories: bulk phenotypes (e.g., area, circularity), global phenotypes (e.g., dry mass density, attenuation density) and local phenotypes (e.g., BF entropy, phase entropy). Previous studies have proven the effectiveness of these features in characterizing different types of cells [1-3] and they were ready for subsequent downstream analysis.

In brief, optical density phenotypes were based on the BF image of the cell whereas mass density phenotypes were extracted from dry mass density map which was converted from the QPI using the well-known linear relationship between refractive index and mass density of most intracellular biomolecules. The slope of this relationship, d*n*/d*c*, is called the specific refractive increment. Specific refractive increments for most biomolecules, (especially those for proteins and nucleic acids) fall within a very narrow range (0.19 ml/g) [4] and thus permits valid evaluation of cell mass inferred from the quantitative phase ($\phi$).

**Bulk features**

Bulk features were extracted according to the mask of the cell in QPI, using basic thresholding. They describe the cell size, cell mass, and the cell shape (i.e. Circularity, Eccentricity, Aspect Ratio, Orientation).

**Global texture features**

In each BF and QPI images, the global texture phenotypes were extracted based on the statistical distribution of the grey-scale values in the images. They include four basic statistical moments of the global distribution (i.e. mean, variance, skewness and kurtosis), and the peak, minimum values and the range of the distribution. Also included is the dry mass density (DMD), which is extracted based on the assumption that the cell in suspension is in spherical shape.

The phase arrangement phenotypes characterize the phase distribution along the radial directions, i.e. distribution of phase times its corresponding radial position. The phase orientation phenotypes on the other hand describe the relationship of phase values, its angular position and angular "repetitiveness". The phase values were first represented in the angular coordinates. Then the distribution was Fourier transformed to obtain a distribution of phase in the angular frequency domain. The statistical moments of this distribution were used as the phenotypes.

We also quantified the centroid displacement and radial distribution of the mass density phenotypes. Centroid displacement measures the displacement of the weighted centroid of the mass/phase from the unweighted centroid obtained from the mask alone. Radial distribution characterizes the tendency of distribution going closer to the edge or to the centre of the cell.

**Local texture features**

To extract the local texture phenotypes, various local filters were used. They include entropy filter with a kernel size of 2μm, standard deviation filter with kernel size of 1 μm, and Hessian-based multiscale filter. The features extracted with these filters have "Entropy", "STD" for optical density features or "DMD contrast" for mass density features, and "Fiber" in their feature names. Hence, DMD Contrasts, QP Entropy and QP Fiber phenotypes are equivalent to BF STD, BF

Entropy and BF Fiber phenotypes, but performed in the quantitative phase map of the cell. We also quantified the centroid displacement and radial distribution of these local texture features.

Finally, the Fit Texture phenotypes were obtained by characterizing the statistical moments of the profile that emphasizes the high spatial frequency of the phase. It was obtained by subtracting the phase profile of the cell with a smoothed phase profile of the cell (computed from a fitted polynomial surface, along the x and y directions up to the 5th degree). The subtracted profile thus contains the high spatial frequency details of the cell.

### Supplementary Note S2: Model architecture of CytoMAD generator

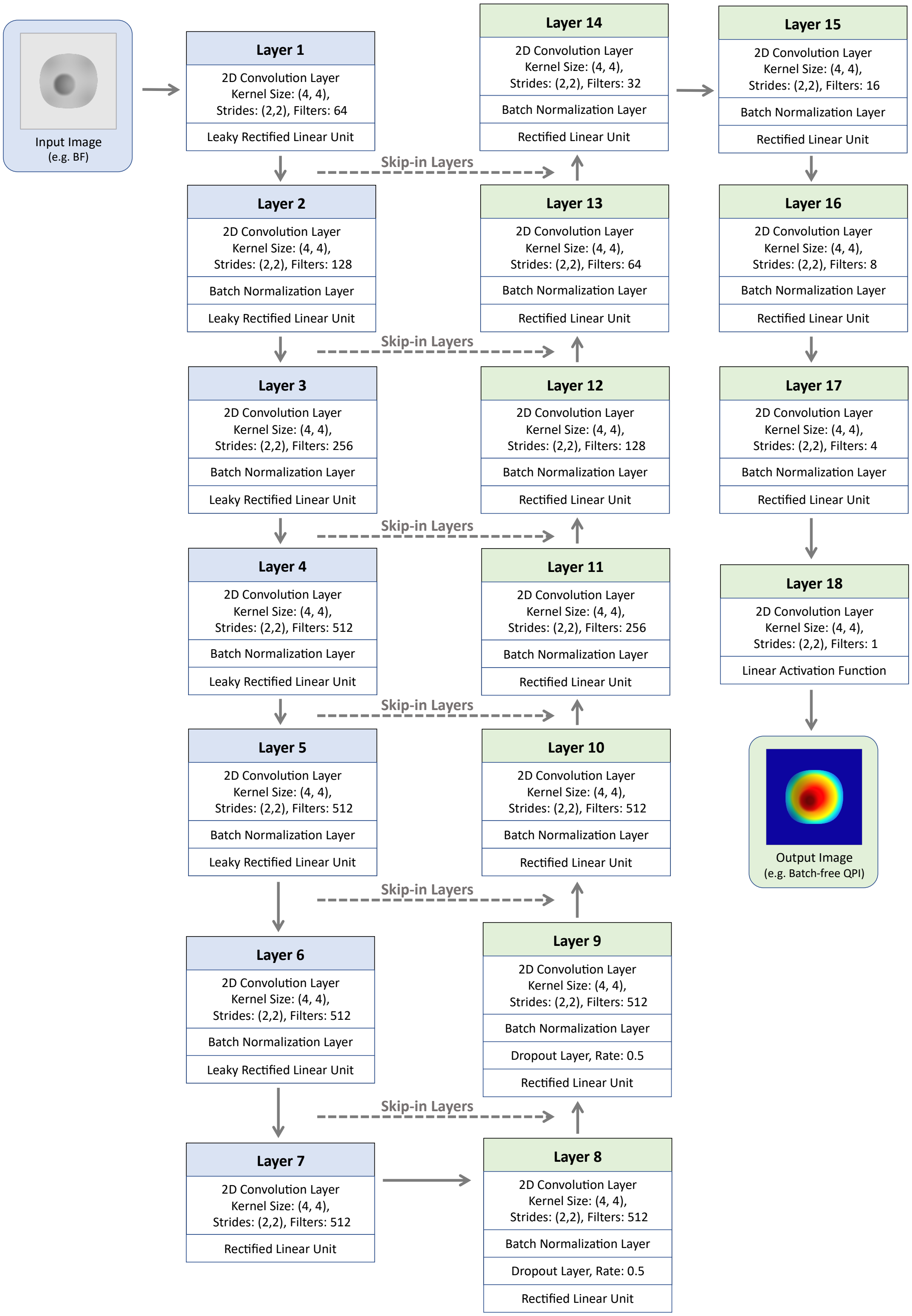

### Supplementary Note S3: Model architecture of classifiers in CytoMAD morphology distillator

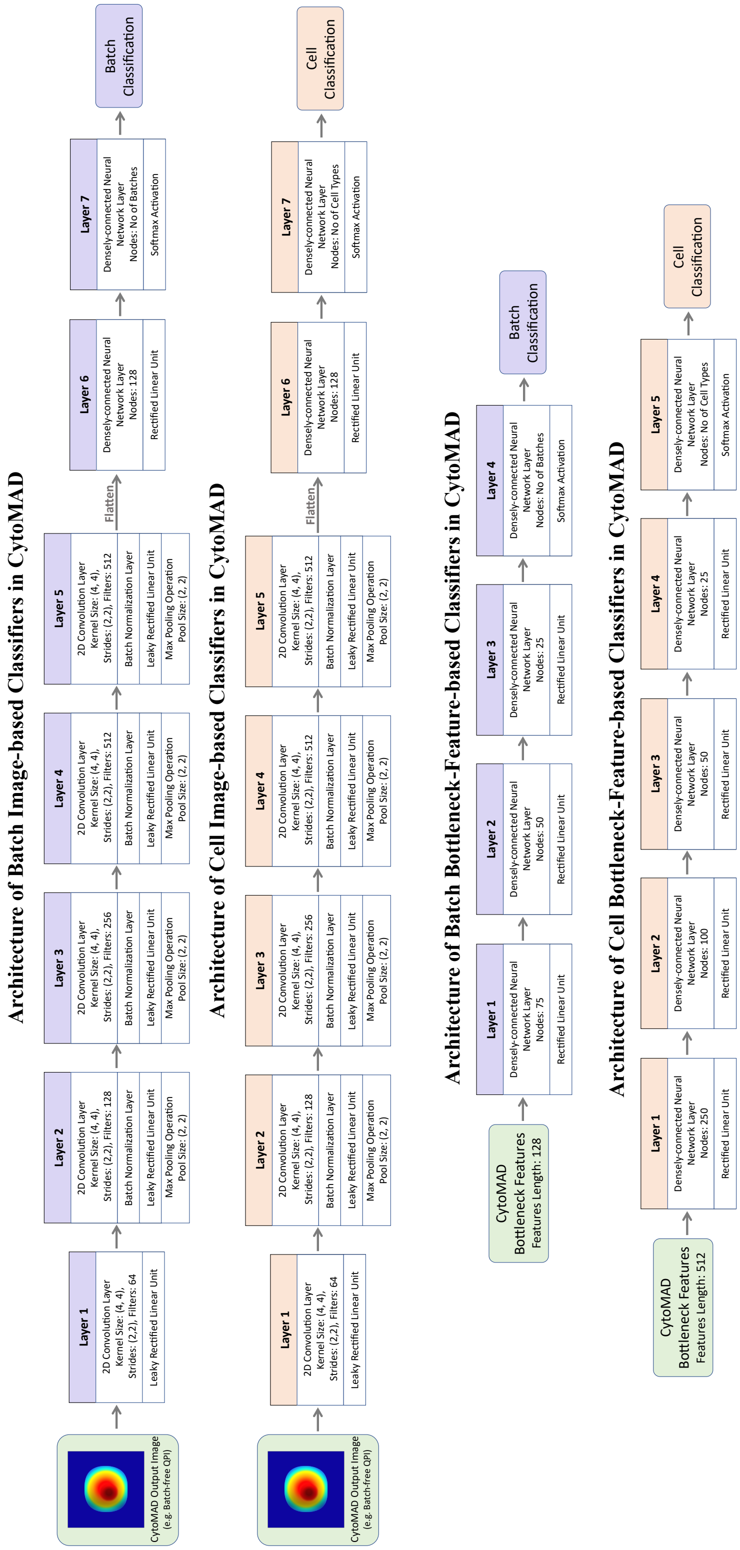

#### Supplementary Note S4: Training details and hyperparameters of CytoMAD

The training of the CytoMAD model was carried out in three main phases. The initial phase involved the pre-training of the cGAN backbone for image contrast translation. This was followed by the pre-training of the morphology distillator, which was a set of classifier networks aiming to discriminate between batch-to-batch variations and biological information in the cell images. The final stage involved the comprehensive training of the CytoMAD model, incorporating the morphology distillator as feedback to guide the generation of batch-distilled phenotypic features (CytoMAD-profile) and cellular images (CytoMAD-images).

##### *Pre-training CytoMAD for Image Contrast Translation*

The cGAN backbone of the CytoMAD model was first trained for basic image-to-image generation and image contrast translation. The model was trained for 100 epochs to convert input images of a particular image contrast (i.e. BF) to a targeted image contrast (i.e. QPI) as output. The learning rate for the cGAN model was set at 0.0002, while the discriminator, providing feedback for the cGAN backbone, was trained at a learning rate of 0.0001. Such pre-training of the CytoMAD produced 1D array at the bottleneck layer, representing cellular phenotypic features (i.e. no-CytoMAD-profile), and output images (i.e. no-CytoMAD-images).

##### *Pre-training of the Morphology Distillator*

With the no-CytoMAD-profile and no-CytoMAD-images from the pre-trained CytoMAD, the morphology distillator was subsequently pre-trained. The morphology distillator is a set of classifier networks which composed of 2 types of classifiers: batch classifiers and cell-type/state classifiers. To identify the batch-to-batch variations and biological information at the bottleneck region of CytoMAD, neural-network-based batch classifiers and cell-type/state classifiers were trained with the no-CytoMAD-profile as input. These classifiers were trained at a learning rate of 0.0001 with 100 epochs. Additionally, convolutional-neural-network-based batch classifiers and cell-type/state classifiers were trained using no-CytoMAD-images with a learning rate of 0.0001 for 100 epochs. With the trained morphology distillator, the model parameters were fixed and the classifiers were implemented at both the bottleneck region and the output of the CytoMAD to provide feedback for guiding the next training round of batch correction.

##### *Comprehensive Batch-distilling Training of the CytoMAD*

In the final training phase, the entire CytoMAD model was trained at a learning rate of 0.0002 for 300 epochs, with the incorporation the morphology distillator as feedback. This comprehensive training enabled the CytoMAD model to generate batch-aware cellular phenotypic features (CytoMAD-profile) and output images (CytoMAD-images).

#### Supplementary Note S5: Training data size of CytoMAD

| | |
|---|---|
| **Lung Cancer Cell Lines** | |
| No. of cell lines/ conditions | 7 types of cell |
| No. of batch | 3 batches |
| Training size | 1,000 cells × 7 types × 3 batches = **21,000 cells** |
| Validation size | 200 cells × 7 types × 3 batches = **4,200 cells** |
| Test size | 40,000 cells × 7 types × 3 batches = **840,000 cells** |
| **Drug Assays of H2170** | |
| No. of cell lines/ conditions | 3 types of drug × 6 different concentration levels = 18 conditions |
| No. of batch | 2 batches |
| Training size | 1,000 cells × 18 conditions × 2 batches = **36,000 cells** |
| Validation size | 500 cells × 18 conditions × 2 batches = **18,000 cells** |
| Test size | 5,000 cells × 18 conditions × 2 batches = **180,000 cells** |
| **Lung Cancer Patients Samples** | |
| No. of cell lines/ conditions | 4 patients,<br>each composed of lung tumors tissue, normal lung tissue and peripheral blood samples |
| No. of batch | 4 batches |
| Training size | 1,000 cells from tumor × 4 batches<br>+ 1,000 cells from blood sample × 4 batches<br>= **8,000 cells** |
| Validation size | 500 cells from tumor × 4 batches<br>+ 500 cells from blood sample × 4 batches<br>= **4,000 cells** |
| Test size | **>120,000 cells** from tumors and peripheral blood,<br>in addition to **~56,000 cells** from unseen patients' normal lung tissue |

#### Supplementary Note S6: Concentration Conditions of Drug Assays of H2170

| Docetaxel | | Afatinib | | Gemcitabine | |
|---|---|---|---|---|---|
| Annotation | Concentration | Annotation | Concentration | Annotation | Concentration |
| **D1** | 0.000376$\mu M$ | **A1** | 0.0019$\mu M$ | **G1** | 0.000751$\mu M$ |
| **D2*** | 0.00376$\mu M$ | **A2*** | 0.019$\mu M$ | **G2*** | 0.00751$\mu M$ |
| **D3** | 0.0188$\mu M$ | **A3** | 0.095$\mu M$ | **G3** | 0.01502$\mu M$ |
| **D4** | 0.0376$\mu M$ | **A4** | 0.19$\mu M$ | **G4** | 0.0751$\mu M$ |
| **D5** | 0.376$\mu M$ | **A5** | 1.9$\mu M$ | **G5** | 0.751$\mu M$ |
| DMSO | | | | | |
| **Control** | 0.367% | **Control** | 0.19% | **Control** | 0.3755% |

* IC50 denotes half-maximal inhibitory concentration of drug.

**Figure S1: Confusion matrix on lung cancer cell types classification with CytoMAD QPI**

**Image-based**

**No CytoMAD**

Predicted Class (columns) / True Class (rows)

| True Class | H69 | H358 | H520 | H526 | H1975 | H2170 | HCC827 |
|---|---|---|---|---|---|---|---|
| H69 | **0.30** | 0.09 | 0.02 | 0.05 | 0.20 | 0.29 | 0.05 |
| H358 | 0.35 | **0.11** | 0.08 | 0.15 | 0.09 | 0.20 | 0.02 |
| H520 | 0.06 | 0.22 | **0.03** | 0.18 | 0.01 | 0.30 | 0.20 |
| H526 | 0.00 | 0.01 | 0.00 | **0.58** | 0.38 | 0.00 | 0.01 |
| H1975 | 0.08 | 0.01 | 0.00 | 0.62 | **0.24** | 0.02 | 0.03 |
| H2170 | 0.09 | 0.09 | 0.11 | 0.10 | 0.02 | **0.53** | 0.04 |
| HCC827 | 0.09 | 0.00 | 0.04 | 0.16 | 0.06 | 0.13 | **0.51** |

**CytoMAD**

Predicted Class (columns) / True Class (rows)

| True Class | H69 | H358 | H520 | H526 | H1975 | H2170 | HCC827 |
|---|---|---|---|---|---|---|---|
| H69 | **0.83** | 0.01 | 0.01 | 0.14 | 0.01 | 0.00 | 0.00 |
| H358 | 0.02 | **0.68** | 0.01 | 0.11 | 0.05 | 0.05 | 0.07 |
| H520 | 0.00 | 0.01 | **0.78** | 0.00 | 0.02 | 0.12 | 0.07 |
| H526 | 0.27 | 0.06 | 0.00 | **0.58** | 0.08 | 0.00 | 0.01 |
| H1975 | 0.01 | 0.00 | 0.01 | 0.03 | **0.93** | 0.01 | 0.00 |
| H2170 | 0.00 | 0.01 | 0.03 | 0.00 | 0.02 | **0.88** | 0.06 |
| HCC827 | 0.02 | 0.09 | 0.05 | 0.01 | 0.01 | 0.07 | **0.75** |

*Left:* Confusion matrix of no-CytoMAD-images. *Right:* Confusion matrix with CytoMAD-images.

**Figure S2: Correlation between CytoMAD-features profile and biophysical phenotypes**

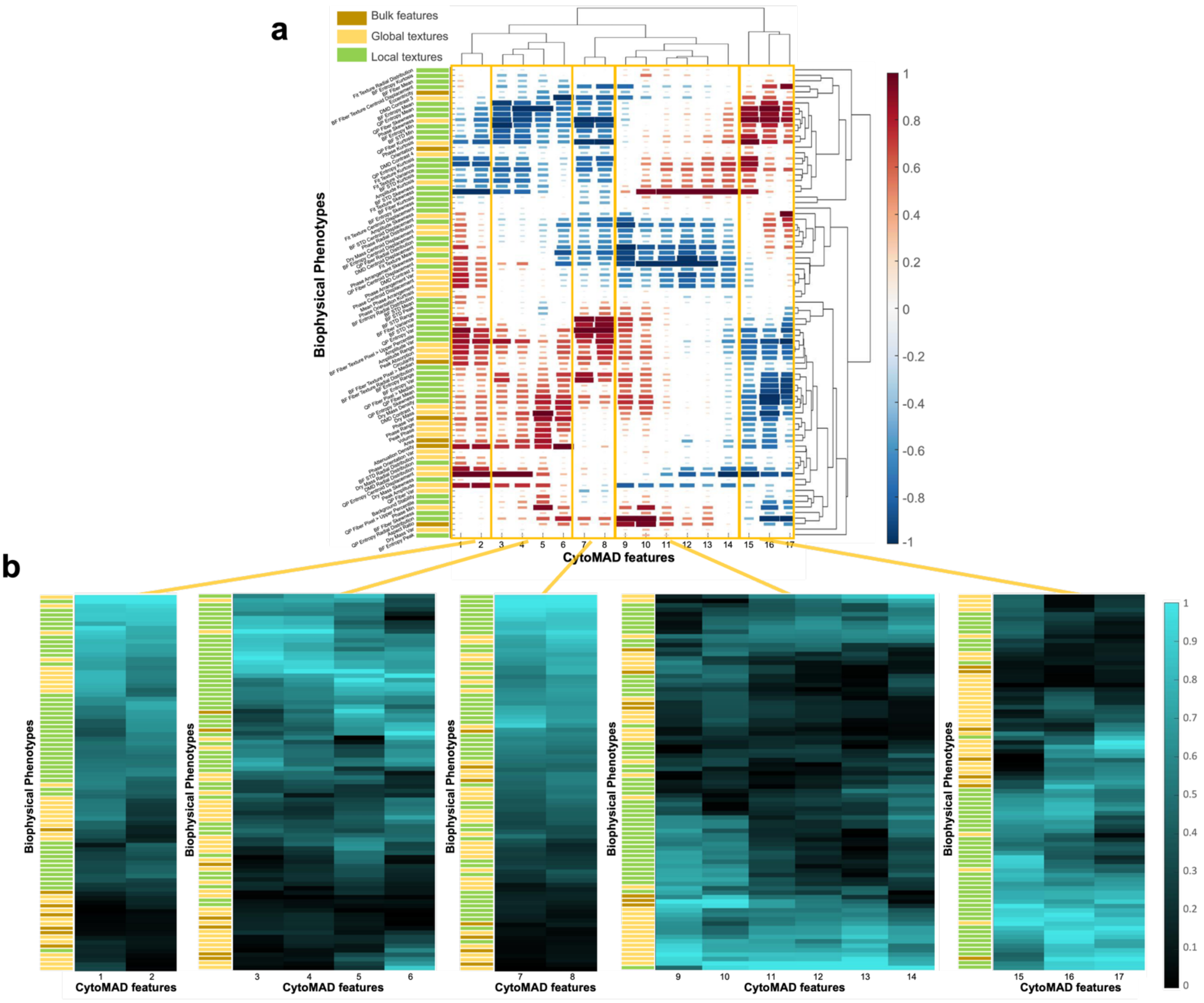


**(a)** Correlation map between CytoMAD-features and biophysical phenotypes. The correlation values are normalized along each CytoMAD feature. Red: positive correlation; Blue: negative correlations. Different colors are also used to represent the 3 categories of the biophysical phenotypes: bulk features (brown), global features (yellow) and local features (green). Hierarchical clustering grouped the CytoMAD-features into 5 groups, as indicated with the rectangles. **(b)** Absolute correlation profile of each CytoMAD-features group. The absolute correlation values are reported.

**Figure S3: CytoMAD-based biophysical phenotypes across major lung cancer types and adenocarcinoma subtypes**

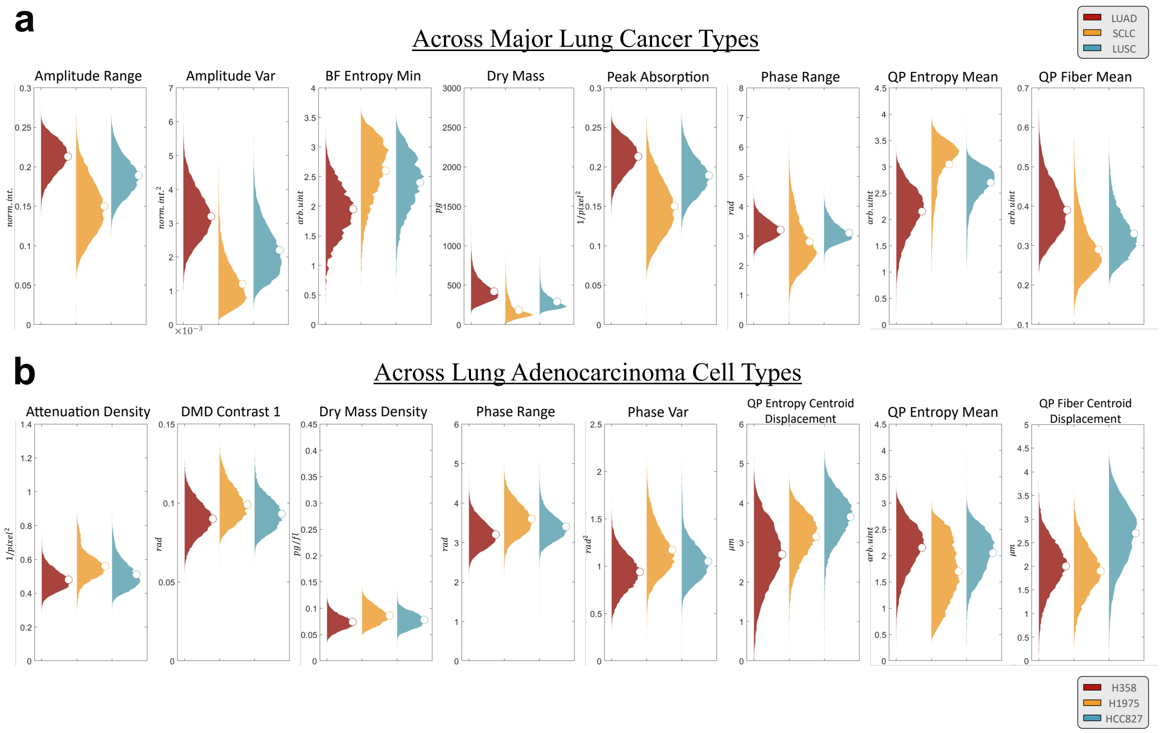


**(a)** Violin plots across major lung cancer types: LUAD denoted in red, SCLC denoted in yellow and LUSC denoted in green. **(b)** Violin plots across lung adenocarcinoma subtypes. Different colors are used to indicate different adenocarcinoma subtypes, with H358 denoted in red, H1975 denoted in yellow and HCC827 denoted in green.

**Figure S4: H2170 drug assays experimental design with CytoMAD**

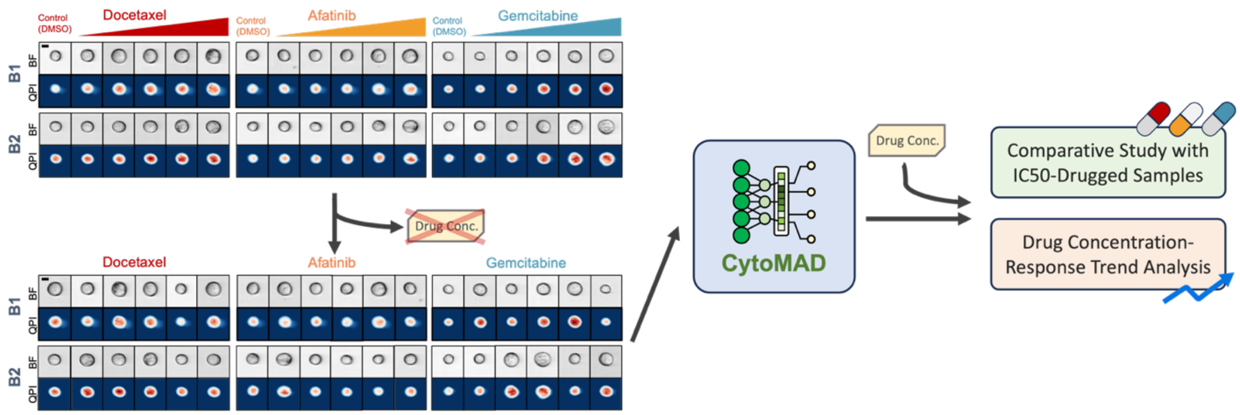


This figure depicts the analysis of H2170 cell drug response in datasets where cells were treated with docetaxel, afatinib, and gemcitabine at varying concentrations across two batches, yielding 18 unique drug treatment conditions (five concentration levels plus one negative control per drug). During the CytoMAD's training phase, the model was only fed batch labels and drug types, with drug concentrations intentionally omitted to test its ability to autonomously discern morphological variations due to concentration differences. Post-training, CytoMAD's outputs facilitated comparative IC50 studies and concentration-response trend analysis for each drug, serving to confirm the model's efficacy in capturing biologically relevant changes and to conduct expansive biological evaluations.

**Figure S5: SSIM and RMSE distributions of H2170 in the drug assays**

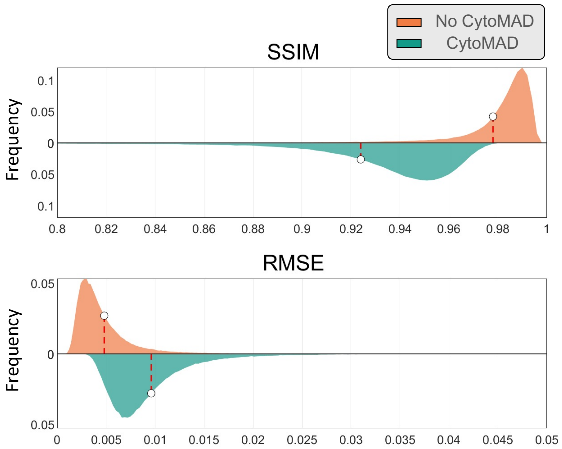


The average structural similarity index (SSIM) for QPI of no-CytoMAD-images and CytoMAD-images were 0.9795 and 0.9241. A high SSIM value denotes a high structural similarity between the generated QPI and the ground truth QPI. While for root mean square error (RMSE), the average values for QPI of no-CytoMAD-images and CytoMAD-images were 0.0049 and 0.0098. A low RMSE value suggests the accurate phase value reconstruction. White circles represent the average values of the distributions.

### Figure S6: Violin plots of biophysical phenotypes across IC50 samples

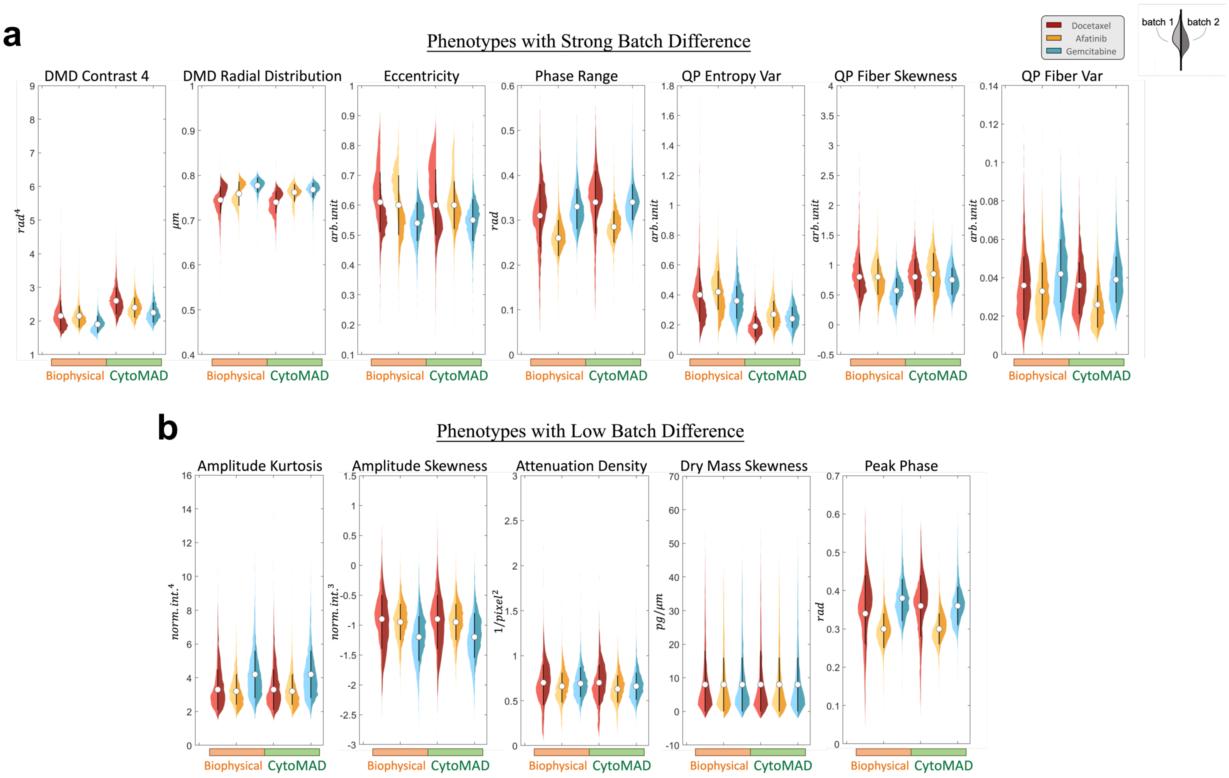


**(a)** Violin plots of biophysical phenotypes of strong batch differences. **(b)** Violin plots of biophysical phenotypes of low batch differences. Different colors are used to indicate different drug treatment samples. The 2 sides of each violins represent 2 different batches and the white circle at the center line each violin denotes the average optophysical phenotype values between the 2 batches. The 3 violins on the left side of each plot indicates the optophysical phenotypic distribution without CytoMAD, while the 3 violins on the right side indicates the optophysical phenotypic distribution with CytoMAD.

**Figure S7: Batch distance correction ratio after CytoMAD**

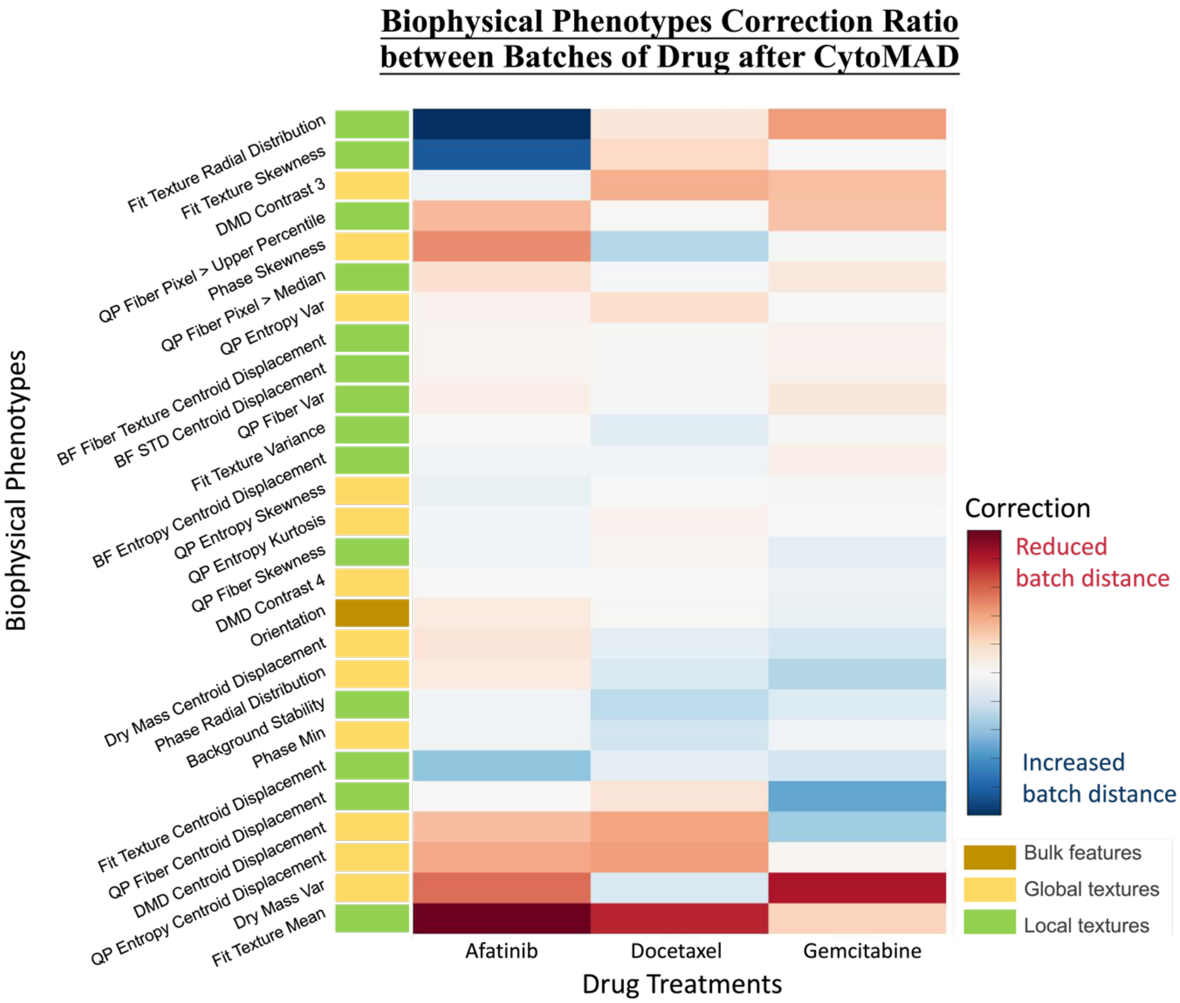


The relative changes in batch distance of each biophysical phenotypes under each drug treatments after implementing CytoMAD were quantified as the correction ratio. The red color denotes a reduction of batch distance after CytoMAD, while the blue color denotes an increase in batch distance after CytoMAD.

**Figure S8: Batch distance on biophysical phenotypes along drug concentration**

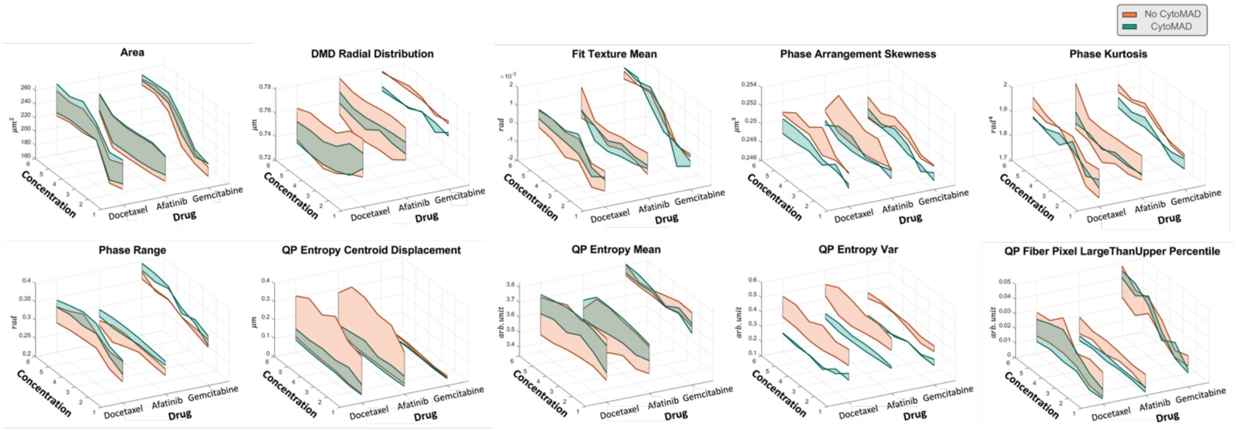


Selected biophysical phenotypes, with their corresponding values in different drug treatments, were plot along the drug concentration. Each orange line on the plot represents a batch of each drug treatments without CytoMAD, and the orange shaded area between lines denotes the phenotypic distance between batches without CytoMAD. While each green line on the plot represents a batch of each drug treatments with CytoMAD, and the green shaded area between lines denotes the phenotypic distance between batches with CytoMAD.

#### Figure S9: NSCLC samples images from multi-ATOM and CytoMAD-images results

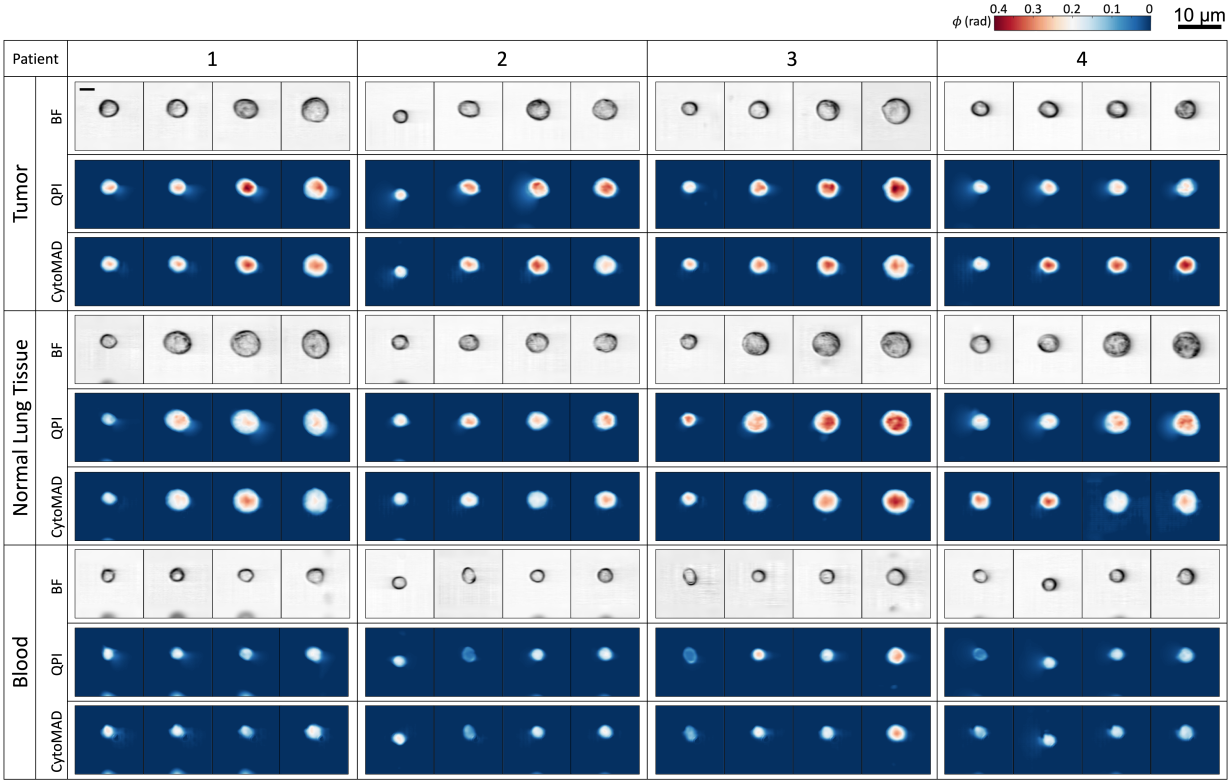


Color bar shows the phase values. Scale-bar: 10μm. FOV: 45μm × 45μm.

#### Figure S10: SSIM and RMSE of NSCLC samples

a

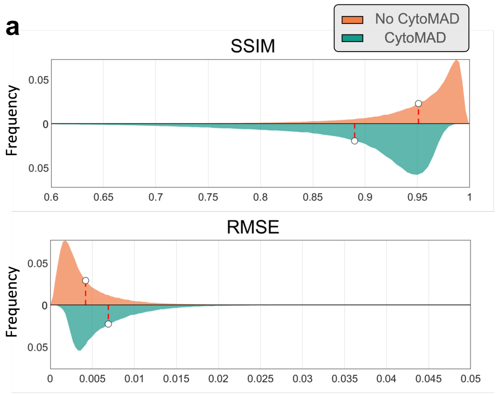


b

**With CytoMAD**

| | SSIM | RMSE |
|---|---|---|
| **Tumor** | 0.9100 | 0.0067 |
| **Normal** | 0.8769 | 0.0126 |
| **Blood** | 0.8774 | 0.0060 |

**(a)** Violin plots on SSIM and RMSE distribution. The average SSIM for QPI of no-CytoMAD-images and CytoMAD-images were 0.9500 and 0.8881. A high SSIM value denotes a high structural similarity between the generated QPI and the ground truth QPI. While for RMSE, the average values for QPI of no-CytoMAD-images and CytoMAD-images were 0.0053 and 0.0084. A low RMSE value suggests the accurate phase value reconstruction. The SSIM and RMSE distributions of no-CytoMAD QPI are indicated in orange, while CytoMAD QPI are indicated in green. The white circles represent the average values of the distributions. **(b)** SSIM and RMSE of different clinical samples. The SSIM and RMSE under various NSCLC samples after implementation of CytoMAD were reported, achieving an overall SSIM > 0.87 and RMSE <0.013 among all samples.

**Figure S11: UMAP on NSCLC blood and tumor samples**

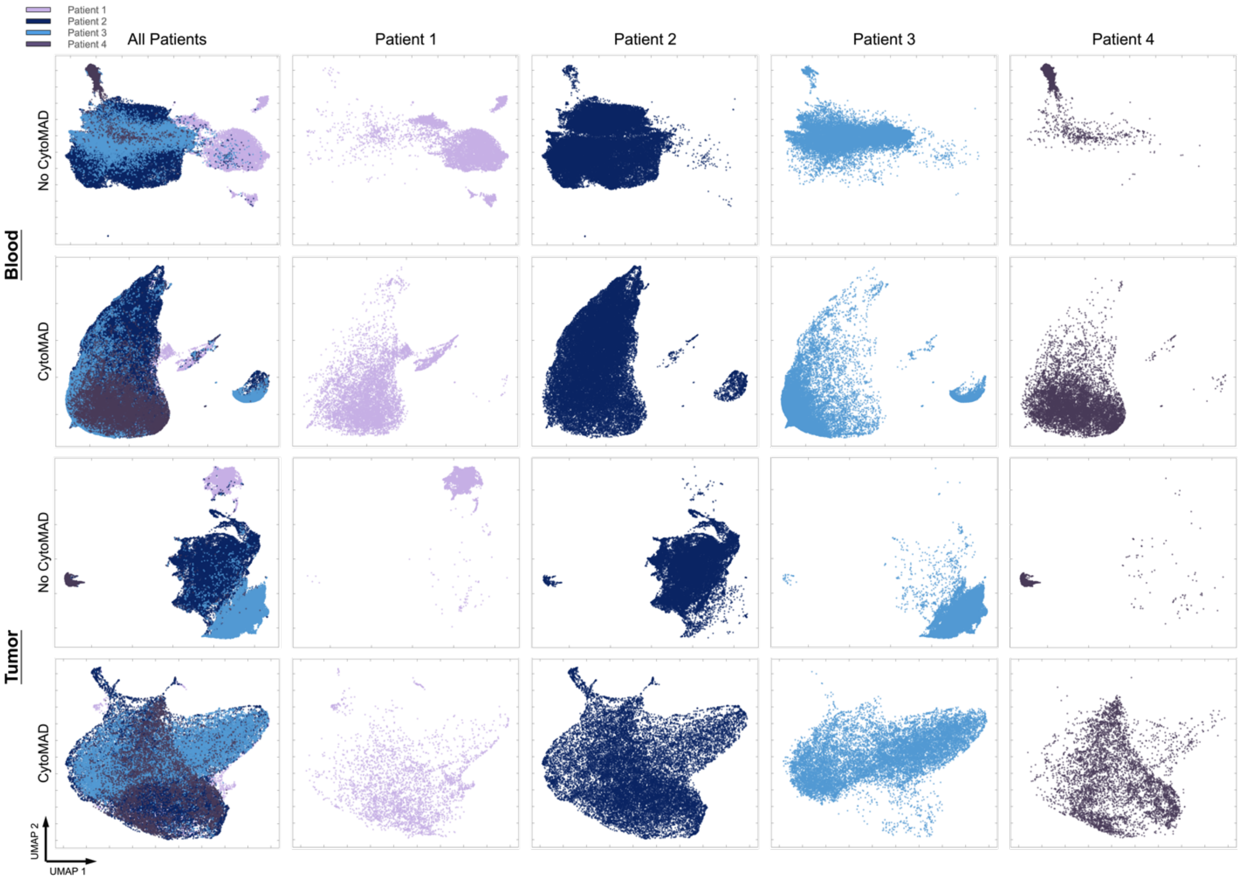


Every data point represents an individual cell. Different colors are used to indicate different batches/patients. To enhance the visualization of overlapping data, individual plots for each patient are provided. *1st-row:* UMAP on patients' blood samples with no-CytoMAD-profile. *2nd-row:* UMAP on patients' blood samples with CytoMAD-profile. 3*rd-row:* UMAP on patients' tumor samples with no-CytoMAD-profile. *4th-row:* UMAP on patients' tumor samples with CytoMAD-profile.

**Figure S12: NSCLC EpCAM and vimentin fluorescence signals and gating**

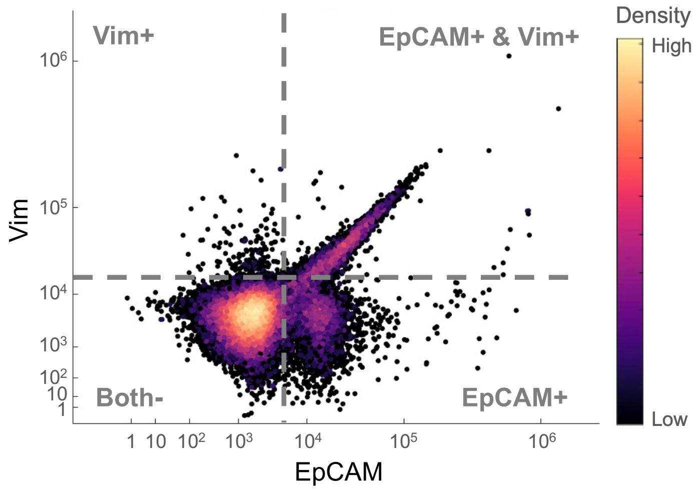


With EpCAM and Vim antibody fluorescence staining, the cells were categorized into 4 groups, including EpCAM positive (EpCAM+), Vim positive (Vim+), both EpCAM and Vim positive (EpCAM+ & Vim+) and non-fluorescence cells (Both-) .Color bar shows the cell density.

**Figure S13: UMAP on different batches/patients of tumor and normal lung tissue samples**

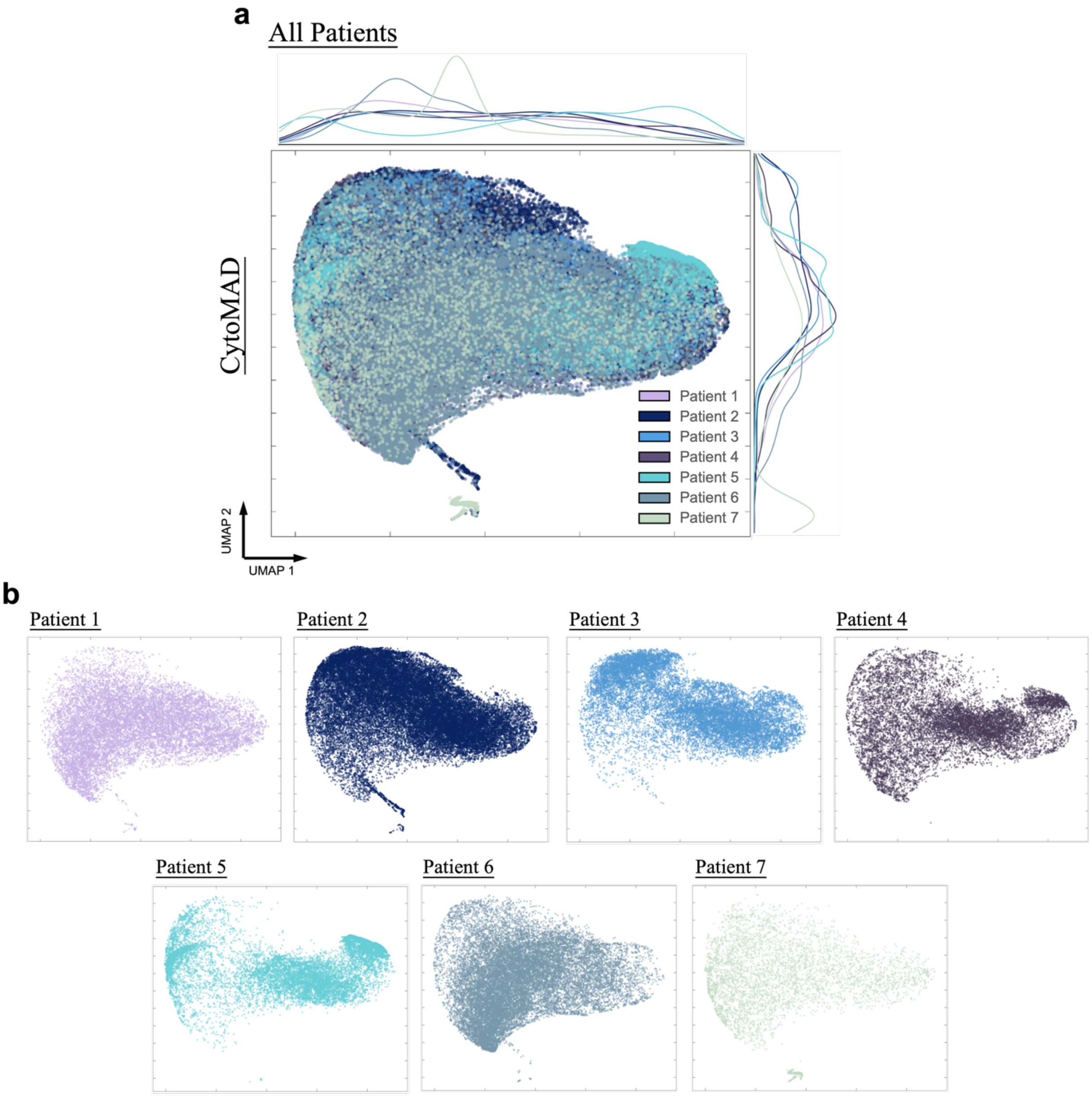


**(a)** UMAP analysis was conducted based on patients' tumors and normal lung tissue samples with CytoMAD-profiles. Different colors are used to indicate different batches/patients. The distributions of each patients are projected on the sides for visualization the highly overlapping populations. **(b)** UMAP of each patient. The 7 patients' data were plotted separately for better visualization of data distribution, with the 3 new, unseen patients denoted as patients 5, 6, and 7 for testing the generalizability of the model across new patients.

**Figure S14: UMAP on molecular markers of tumor and normal lung tissue samples**

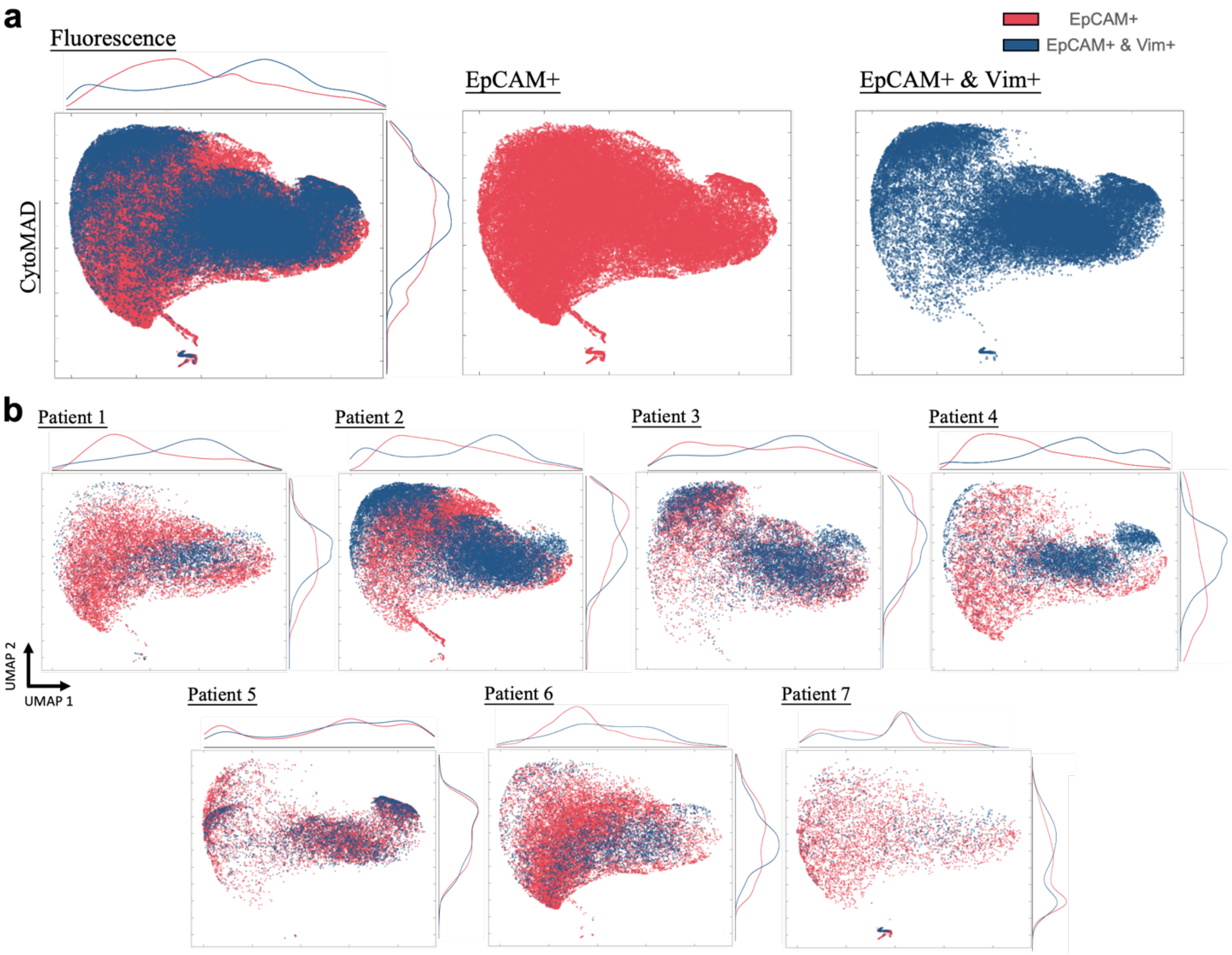


**(a)** UMAP analysis was conducted based on patients' tumors and normal lung tissue samples with CytoMAD-profiles. Different colors are used to indicate different molecular markers of EpCAM+ and Both+ (i.e. EpCAM+ & Vim+). The distributions of each molecular markers are projected on the sides for visualization the highly overlapping populations. Different molecular markers' data were plotted separately for better visualization of data distribution. **(b)** UMAP of each patient. The 7 patients' data were plotted separately for better visualization of data distribution.

**Figure S15: Z-Score of subpopulation analysis in EpCAM+Vim+ cells**

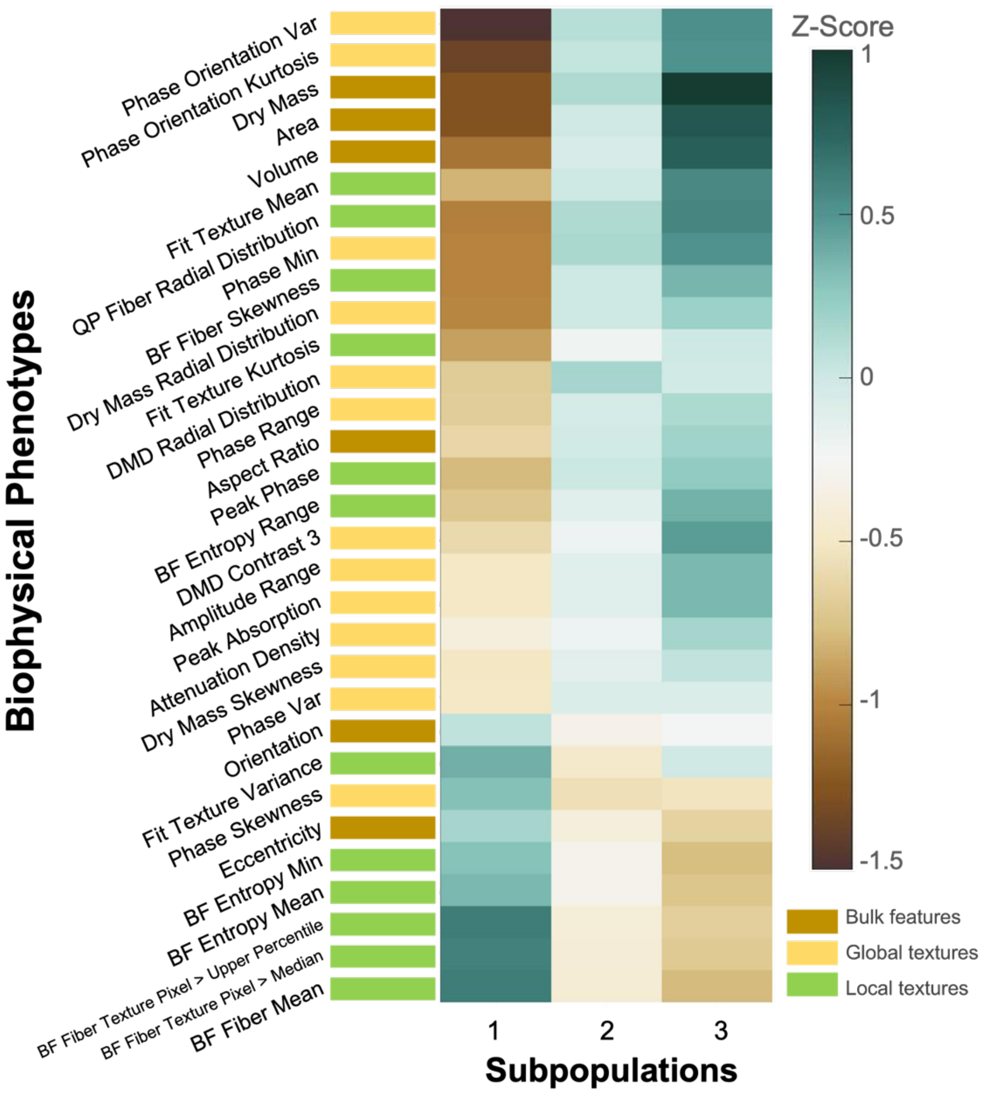


The z-score of the subpopulations are reported, with green color denotes a positive z-score and brown denotes a negative z-score Different colors are used to represent the 3 categories of the biophysical phenotypes.

**Figure S16: UMAP on intra-batch analysis**

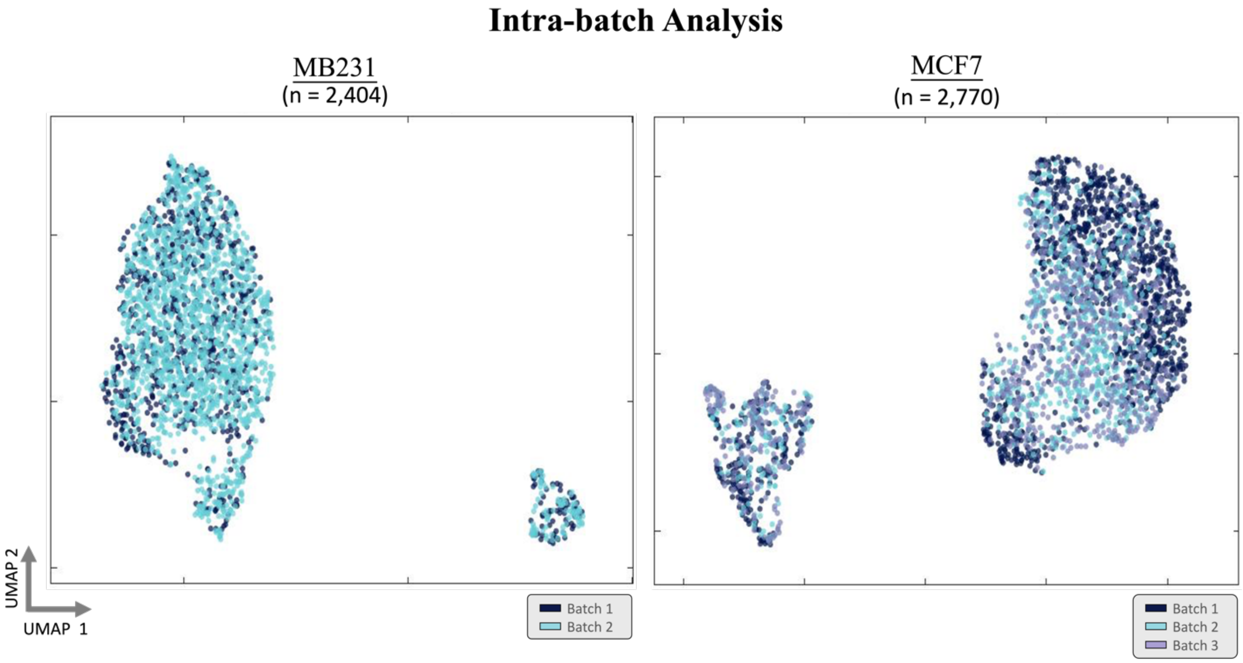


Comparative Analysis of Intra-Batch Variations in Flow Imaging Cytometry. A dedicated investigation was conducted on another flow imaging cytometry technique [5-7], with a specific focus on breast cancer cell types, MDA-MB231, and MCF7. Multiple batches for each cell type were acquired on the same day under consistent optical system settings but at different times. UMAP analysis revealed no observable differences within the same batches, suggesting that intra-batch variations in this dataset can be considered negligible.

#### Figure S17: UMAP on H520 cell samples with CytoMAD

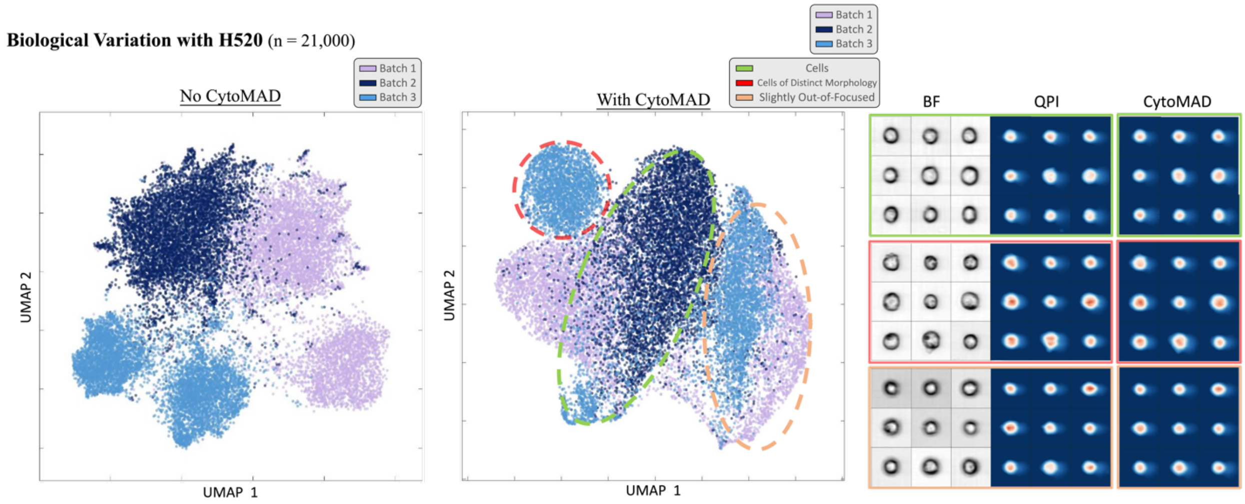


With 3 batches of H520 cell samples from LUSC, UMAP analysis was conducted to investigate CytoMAD's impact on batch-to-batch variations. Applying CytoMAD, while the batches appeared more cohesive (middle), we could still discern three clusters indicative of subtle cellular condition differences—normal cells, cells with distinct morphology, and some residual cells which are slightly out-of-focus. This visual comparison confirms CytoMAD's ability to preserve subtle intra-batch heterogeneity, ensuring biological traits are retained while seamlessly reducing inter-batch variability. Thus, CytoMAD adeptly minimizes batch effects, enhancing data integration (e.g., the significant improvement in classification accuracy in Fig. 2c) without indiscriminately conflating batches

**Figure S18: Benchmarking with existing batch-correction algorithms**

**a**

| | | HARMONY | MNN | CytoMAD |
|---|---|---|---|---|
| Minimizing Batch Effect | 1-dimensional Data | ✓ | ✓ | ✓ |
| | 2-dimensional Data (i.e. Images) | ✗ | ✗ | ✓ |
| Data Interpretability | | ✗ | ✗ | ✓ |

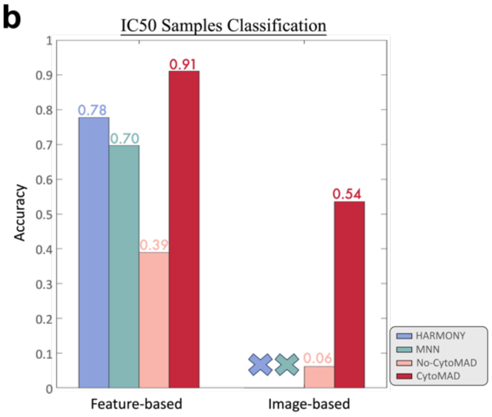


**(a)** This table provides a comparative overview of CytoMAD with prevalent batch effect removal tools such as HARMONY and MNN, which are traditionally used in single-cell omics. It underscores the inherent limitations of these established methods when confronted with 2D imaging data, which is CytoMAD's domain of operation. The table also notes that while many current algorithms employ dimensionality reduction to achieve stable batch correction in single-cell RNA sequencing, this often results in a loss of interpretability, particularly in the context of cellular morphology. In contrast, CytoMAD is specifically designed to take 2D images as inputs, offering outputs in the form of batch-corrected 1D feature profiles and 2D images. This preserves the granular detail necessary for comprehensive biological analysis and enables a more nuanced understanding of biological phenomena.

**(b)** This section details a benchmarking study comparing CytoMAD with HARMONY, MNN, and a baseline cGAN model using IC50 samples from the drug assays of the H2170 dataset. The evaluation of batch removal efficacy involved training NN-based and CNN-based classifiers on CytoMAD-profile and CytoMAD-images, respectively. The resulting classification performance across various algorithms was reported for both feature-based and image-based classifications. CytoMAD demonstrated superior performance in feature-based classification, achieving an across-batch accuracy of 0.91, surpassing HARMONY (0.78) and MNN (0.70). The model showcased a significant improvement over the basic cGAN model (0.39 accuracy), highlighting its efficacy in overcoming batch-to-batch variations. In image-based classification, CytoMAD outperformed, achieving a substantially higher accuracy of 0.54 compared to 0.06 without CytoMAD. Notably, as HARMONY and MNN are designed for 1D data, they were not applicable in this aspect of the benchmarking. Overall, these results underscore the effectiveness of CytoMAD in mitigating batch effects, affirming its potential as a superior alternative to existing batch effect correction methods *in the context of 2D imaging data.*

**Figure S19: Pre-processing pipeline of multi-ATOM images**

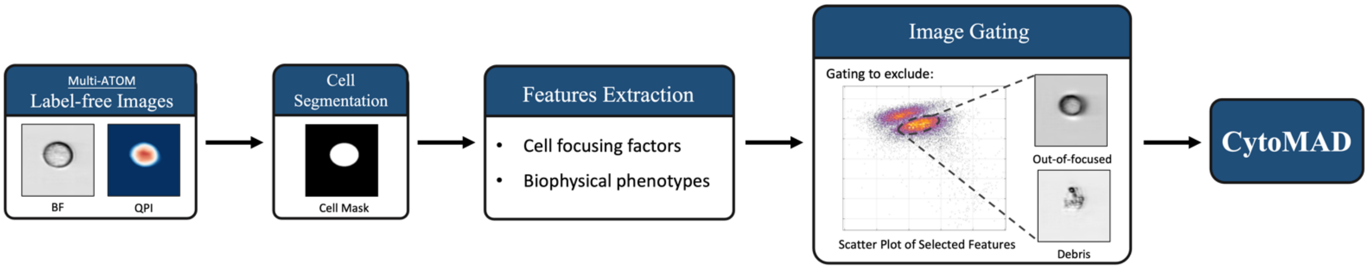


The pre-processing pipeline for multi-ATOM images involved initial cell segmentation on both bright-field (BF) and quantitative phase imaging (QPI) images to distinguish cell bodies from the background. Subsequently, a comprehensive set of features, encompassing cell focusing factors and biophysical phenotypes, was quantified based on the identified cell masks. A specific subset of cell features was chosen to create 2-dimensional scatter plots, facilitating image inspection and cell gating. This gating process effectively excluded undesirable cells, such as those that were out-of-focus or debris, from the datasets. The gated data were then utilized in the training and testing phases of the CytoMAD model.
